# Parallel Evolution of a Drimenol Biosynthetic Gene Cluster Unique to Marine Bacteria

**DOI:** 10.1101/2025.09.20.677545

**Authors:** Nhu Ngoc Quynh Vo, Yuhta Nomura, Shunji Takahashi

**Author notes:** Plant Genomic Network Research Team, RIKEN Center for Sustainable Resource Science, 1–7–22 Suehiro-cho, Yokohama, Kanagawa 230–0045, Japan (Nhu Ngoc Quynh Vo).

## Abstract

Natural bacterial communities represent a rich resource for the discovery of enzymes and natural compounds including terpenoids. However, the biosynthetic potential of marine bacteria remains poorly explored due to limitations in studying fastidious microorganisms. In a recent study, we discovered unusual haloacid dehalogenase (HAD)-like drimenol synthases (DMSs) of marine bacterial origin, including *Aquimarina spongiae* AsDMS and *Flavivirga eckloniae* FeDMS, which produce drimenol *in vitro*. In the present study, we conducted an *in vivo* investigation of the biosynthesis of this drimane-type sesquiterpene in marine bacteria. To verify drimenol accumulation in cultures of *F. eckloniae*, we mined the drimenol biosynthetic genes encoded in the *F. eckloniae* genome and found a putative AraC family transcriptional regulator in the drimenol gene cluster. Upon the addition of arabinose, we observed drimenol biosynthesis in *F. eckloniae*, demonstrating the importance of genomics-driven strategies in investigating natural products. Syntenic comparisons of drimenol gene clusters across bacteria revealed the unique evolution of drimenol biosynthesis in marine bacteria. Moreover, the distinct nature of drimenol pathways and relevant enzymes in bacteria compared to those in plants and fungi suggests parallel evolution. Remarkably, we describe two individual proteins from *Sorangium cellulosum*, HAD and terpene synthase with β domain architecture, similar to HAD-like and terpene synthase β domains of AsDMS, respectively, suggesting the fusion of monodomain proteins during the evolution of bacterial bifunctional DMSs. Our findings shed light on the evolutionary trajectory of the drimenol gene cluster unique to marine bacteria, while also demonstrating the new bacterial mono-β-domain sesquiterpene synthase.

**ABSTRACT GRAPHIC:** 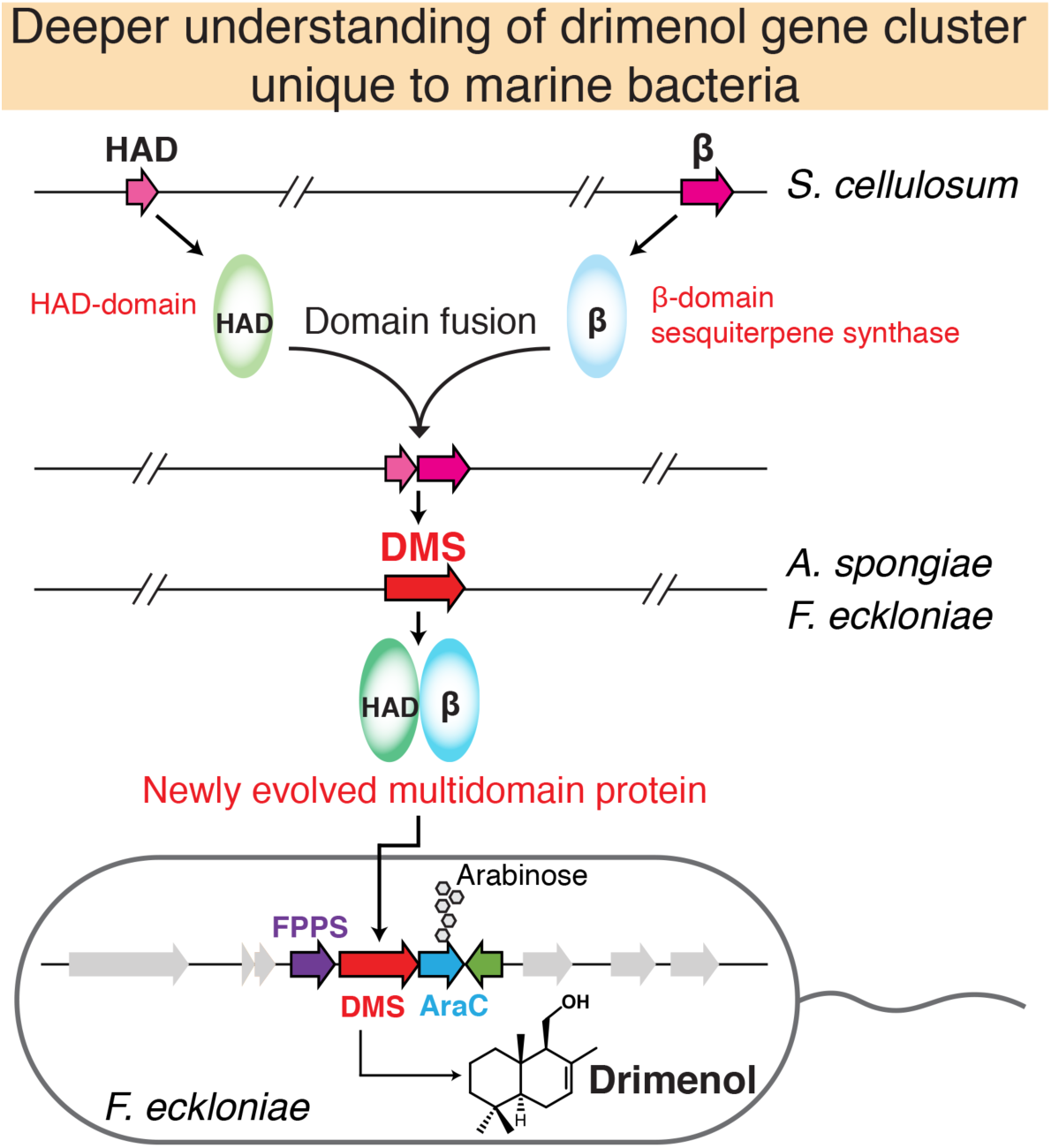

## INTRODUCTION

Natural products derived from bacteria represent an indispensable source of bioactive agents, with more than 50,000 such agents revealed to date.^1–2^ In natural environments, such molecules confer bacteria with a diverse array of ecological and physiological functions ranging from communication to competition.^3–4^ These natural products include examples widely used for therapeutic and biotechnological applications.^5–6^ As the largest class of natural products, terpenoids exhibit a broad spectrum of biological and pharmacological activities.^7–9^ Members of this class of compounds have mainly been identified through studying cultivable bacteria; however, the vast majority of microbes, including those inhabiting the open ocean, have not yet been cultivated.^6^ This cultivation bias has limited the exploration of new bioactive compounds and their genetically encoded production pathways to bacteria, particularly those in marine environments.

In a previous study, we discovered the first examples of bifunctional haloacid dehalogenase (HAD)-like drimenol synthases (DMSs) of marine bacterial origin through bacterial genomic data analysis and *in vitro* enzymatic assays.^10^ These enzymes use farnesyl pyrophosphate (FPP), a linear prenyl diphosphate precursor, as a substrate to produce drimenol, a strong broad-spectrum antifungal agent used as the central biosynthetic precursor for natural drimane sesquiterpenes, a family of bioactive C_15_ terpenoids. The biosynthetic pathways of drimenol and its derivatives have been elucidated in plants^11–12^ and fungi;^13–16^ they involve several oxidases, particularly cytochrome P450 monooxygenases (P450s), in addition to FPP synthase (FPPS) and DMS. However, drimenol biosynthesis remains unknown in marine bacteria. As we previously observed the involvement of marine bacterial DMSs in drimenol biosynthesis *in vitro*, we questioned whether marine bacterial DMSs are truly involved in drimenol biosynthesis in marine bacteria *in vivo*, and whether marine bacteria could produce drimenol under laboratory conditions, which would elucidate the roles of ocean bacteria in secondary metabolite production. Moreover, the DMS characterized from *Aquimarina spongiae* (AsDMS) in our previous study^10^ exhibited architectural fusion of the HAD-like domain (N-domain) and terpene synthase β domain (C-domain), which uniquely differentiated AsDMS from other known class I terpene synthases that feature α, αβ, or αβγ architectures and class II enzymes that exhibit βγ or αβγ architectures.^17–19^ A single β-domain terpene synthase has been described only for a merosterolic acid synthase that exhibits meroditerpene cyclization activity.^20^ Although understanding the different enzyme origins of each domain of HAD-like sesquiterpene synthases such as AsDMS and their subsequent unique fusion is intriguing, the evolutionary processes driving this protein architecture remain unknown. Furthermore, no studies have examined the evolution of drimenol biosynthetic gene clusters among the biological domains, including bacteria.

Here, we used *Flavivirga eckloniae*, which contains the *FeDMS* gene, as a representative species to investigate drimenol biosynthesis *in vivo* in marine bacteria. Since we were unable to observe drimenol in bacterial cultures, we mined core drimenol biosynthetic genes encoded in the *F*. *eckloniae* genome and found a cluster of AraC family transcriptional regulator with the *FPPS* and *DMS* genes. From these, we treated *F*. *eckloniae* cultures with arabinose and verified drimenol accumulation upon arabinose addition. Syntenic comparisons of gene cluster content and arrangement in terrestrial and marine bacterial genomes indicated the unique evolution of drimenol biosynthesis in marine bacteria, which was found to have evolved in parallel to that in plants and fungi. Strikingly, our thorough sequence and structural model analysis and biochemical validation demonstrated that the bacterial origin-like enzymes of each HAD-like and terpene synthase β domain were monodomain proteins that may have fused to form HAD-like DMSs, including AsDMS, thereby revealing the evolutionary trajectory of a drimenol gene cluster specific to marine bacteria.

## RESULTS AND DISCUSSION

### Metabolic Profiling of Drimenol-type Sesquiterpenes in *F*. *eckloniae*

To investigate whether marine bacteria can produce drimenol and its derivatives *in vivo* under laboratory conditions, we cultivated the commercially available *F*. *eckloniae* ECD14T strain, which harbors the *FeDMS* gene. After fermenting *F*. *eckloniae* in marine broth medium overlaid with dodecane at 25 °C for 2 days, we were unable to observe the production of any drimenol-type sesquiterpenes in the culture and dodecane layer, although (+)-nootkatone (internal standard) and authentic drimenol in the positive control were detected (Supplementary Figure S1). This result may be attributable to the fact that *FeDMS* was not constitutively expressed under our cultivation conditions; conceivably, it is expressed sufficiently only under limited conditions that require chemical inducers, as proposed for other sesquiterpene synthase genes involved in secondary metabolism.^21–22^ This observation suggests the involvement of transcriptional regulators in drimenol biosynthesis in marine bacteria.

### Identification of the Potential Gene Clusters Essential for Drimenol Synthesis in Marine Bacteria

The biosynthesis of drimenol-type sesquiterpenes requires FPPSs responsible for the synthesis of FPP and DMSs for the conversion of FPP to drimenol. Furthermore, the transcriptional regulators may also exist, as described above. Therefore, we investigated the possibility that such clustered key genes are responsible for drimenol production in *F*. *eckloniae* and that their expression may be activated by regulatory genes. We searched the genomic regions surrounding *FeDMS* using the National Center for Biotechnology Information (NCBI) RefSeq Genome Database and found three putative genes closely co-localized with *FeDMS*: an *FPPS* gene (PFAM PF00348; InterPro IPR000092), an AraC family transcriptional regulator (*AraC*-like; IPR020449), and an antibiotic biosynthesis monooxygenase (*ABM*-like; IPR011008) (Figure 1). This gene clustering was highly conserved in other marine bacterial genomes, including *A*. *spongiae*, *Aquimarina* sp. AU474, *Flavivirga amylovorans*, *Flagellimonas* sp. HSM57, and *Aquimarina* sp. AU119. In contrast, the *Rhodobacteraceae* KLH11 lacks the clustered *AraC*-like transcriptional regulator and *ABM*-like gene.

**Figure 1.**
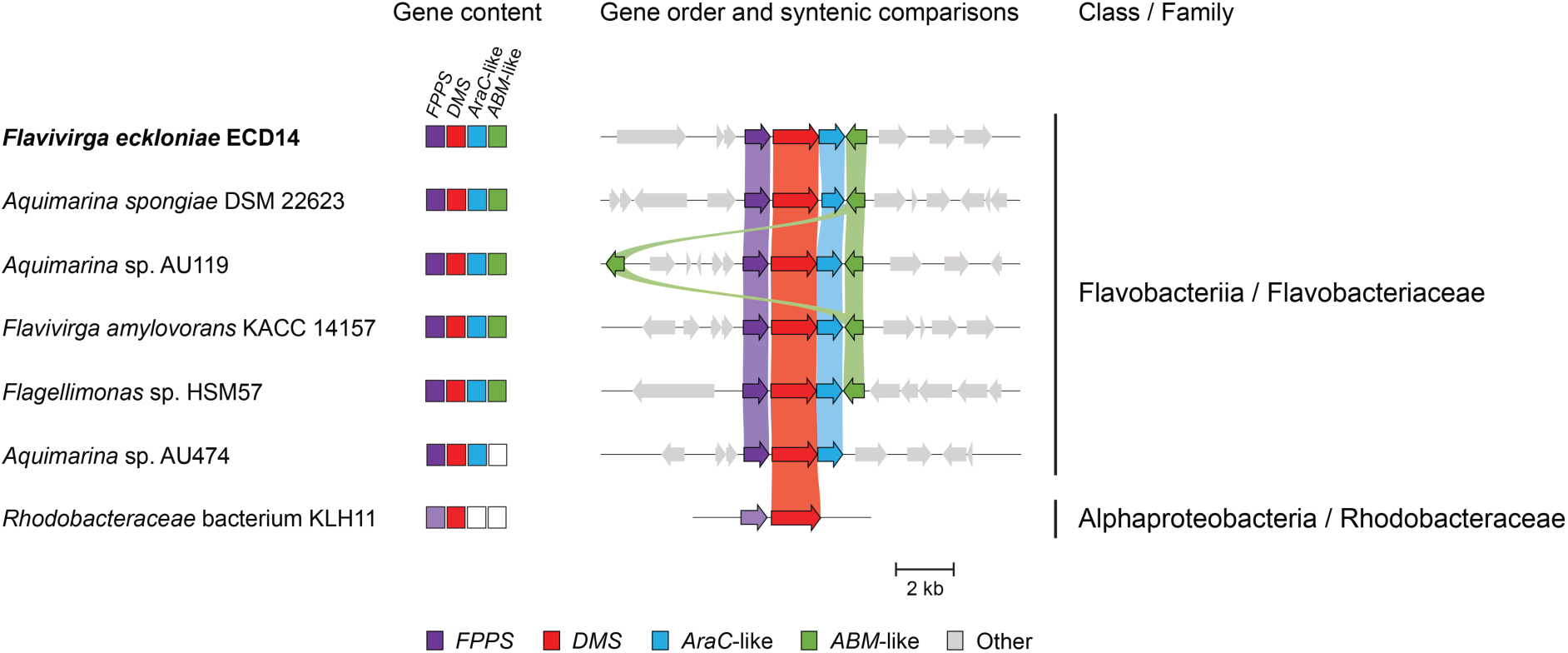
Syntenic comparisons of possible drimenol gene clusters in *Flavivirga eckloniae* (bold black font) and other marine bacteria, with class and family indicated. Central panel shows the presence (colored squares) or absence (white squares) of putative cluster genes encoding a farnesyl pyrophosphate synthase (*FPPS*), a drimenol synthase (*DMS*), a AraC family transcriptional regulator (*AraC*-like), and an antibiotic biosynthesis monooxygenase (*ABM*-like). Right panel shows the synteny of drimenol cluster organization. The *AraC*-like and *ABM*-like genes were absent from the *Rhodobacteraceae* KLH11 bacterial genome. Genes encoding proteins of similar putative functions with amino acid sequence identity > 30 % are connected by a ribbon. The putative *Rhodobacteraceae* FPPS (sequence identity < 30 %) is shown in light purple. Genes not present in the drimenol cluster are shown in gray.

Phylogenetic and sequence comparison analyses of these putative FPPSs and ABMs showed that they share close evolutionary relationships and typical conserved motifs with functionally characterized and unknown FPPSs and ABMs, respectively (Supplementary Figures S2–S4). The putative AraC family transcriptional regulator proteins contain a conserved C-terminal helix-turn-helix DNA binding domain and share a high degree of homology with homologs from other bacteria (Supplementary Figures S5, S6). The enzymatic activities of DMSs were confirmed in our previous study,^10^ and those of FPPSs as FPP synthases were confirmed in the present study. Accordingly, *in vitro* enzymatic activity assays of FPPSs in the presence of the substrates isopentenyl pyrophosphate (IPP) and dimethylallyl pyrophosphate (DMAPP) or geranyl pyrophosphate (GPP) as an allylic substrate with alkaline phosphatase (BAP) treatment yielded a single compound corresponding to authentic farnesol in terms of its retention time (16.13 min) and exact mass fragmentation pattern (parent mass *m/z* = 222) (Supplementary Figure S7). Interestingly, when we further examined the coupling activity of FPPS with DMS proteins through *in vitro* enzymatic activity assays of these coupled proteins using IPP and DMAPP or GPP as substrates, with or without BAP treatment, in all cases, a single compound was found to correspond to authentic (–)-drimenol (drim-7-en-11-ol) in terms of its retention time (16.86 min) and exact mass fragmentation pattern (parent mass *m/z* = 222) (Figure 2A, B). Similarly, when DMAPP was used as a counter substrate, DMS activity was reduced compared to that observed using GPP, due to the low levels of FPP produced (Figure 2C). Despite our extensive analysis of *in vitro* biochemical assays of ABM enzymes with FPP, farnesol, and drimenol as substrates, we were unable to detect any evident products (Method S1 and Supplementary Figure S8).

**Figure 2.**
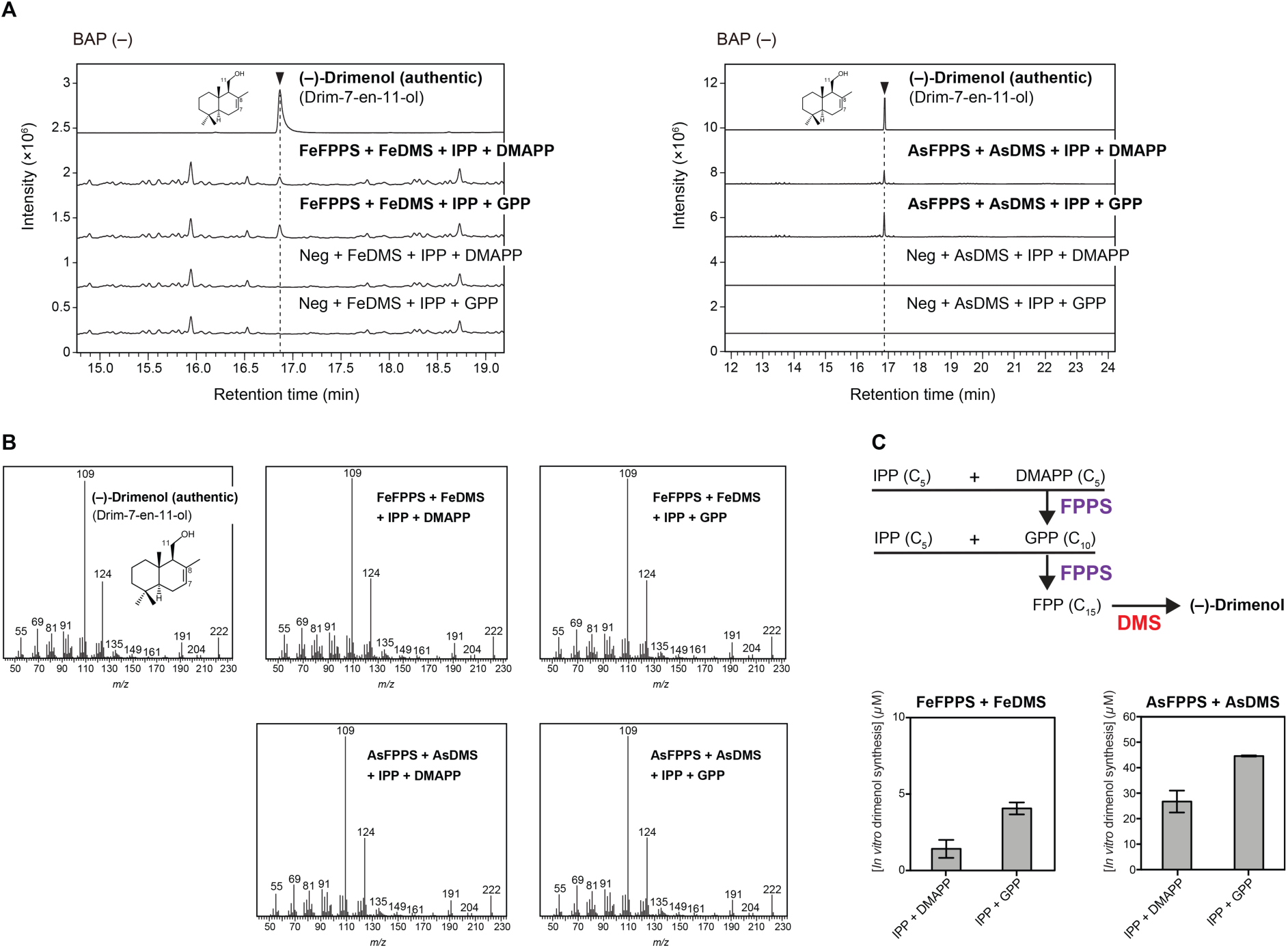
Coupling catalytic activity of the recombinant FeFPPS and AsFPPS proteins with drimenol synthases (DMSs). (A) Gas chromatography–mass spectrometry (GC-MS) chromatograms of the authentic (–)-drimenol (drim-7-en-11-ol) and *in vitro* enzyme reactions of FeFPPS coupled with FeDMS (left panel) or AsFPPS with AsDMS proteins (right panel) using the substrates isopentenyl pyrophosphate (IPP) and dimethylallyl pyrophosphate (DMAPP) or geranyl pyrophosphate (GPP). Reactions were performed without alkaline phosphatase (BAP). (B) Mass spectra of authentic drimenol and reaction products detected in (A). The GC-MS data in (A) represent three independent reactions performed with the same preparation of purified protein. (C) Reaction pathway (upper panel) and comparison of drimenol production by an *in vitro* coupling assay of FPPS with DMS proteins (lower panel). Data are means ± standard deviation (SD) of three independent experiments. Reactions in the absence of FPPS proteins were used as negative controls (Neg). Fe, *Flavivirga eckloniae*; As, *Aquimarina spongiae*. *m/z*, mass-to-charge ratio.

### Detection of Drimenol in Marine Bacteria Cultivated with L-Arabinose Supplementation

We were unable to detect (–)-drimenol in *F*. *eckloniae* under laboratory cultivation without induction. Based on observations of the putative AraC family transcriptional regulator in the drimenol gene cluster, we supplemented the *F*. *eckloniae* culture with 0.02%, 0.2%, or 2% L-arabinose, followed by metabolite extraction and gas chromatography and mass spectrometry (GC-MS) analysis. The results showed that (–)-drimenol was detected only in *F*. *eckloniae* cultured with 0.2% L-arabinose (Figure 3); no (–)-drimenol was detected in the non-supplemented control. These results suggest that L-arabinose plays a role as an activating sugar inducer for the expression of the drimenol gene cluster, and thus the potential involvement of the AraC-family transcriptional regulator in drimenol biosynthesis in marine bacteria.

**Figure 3.**
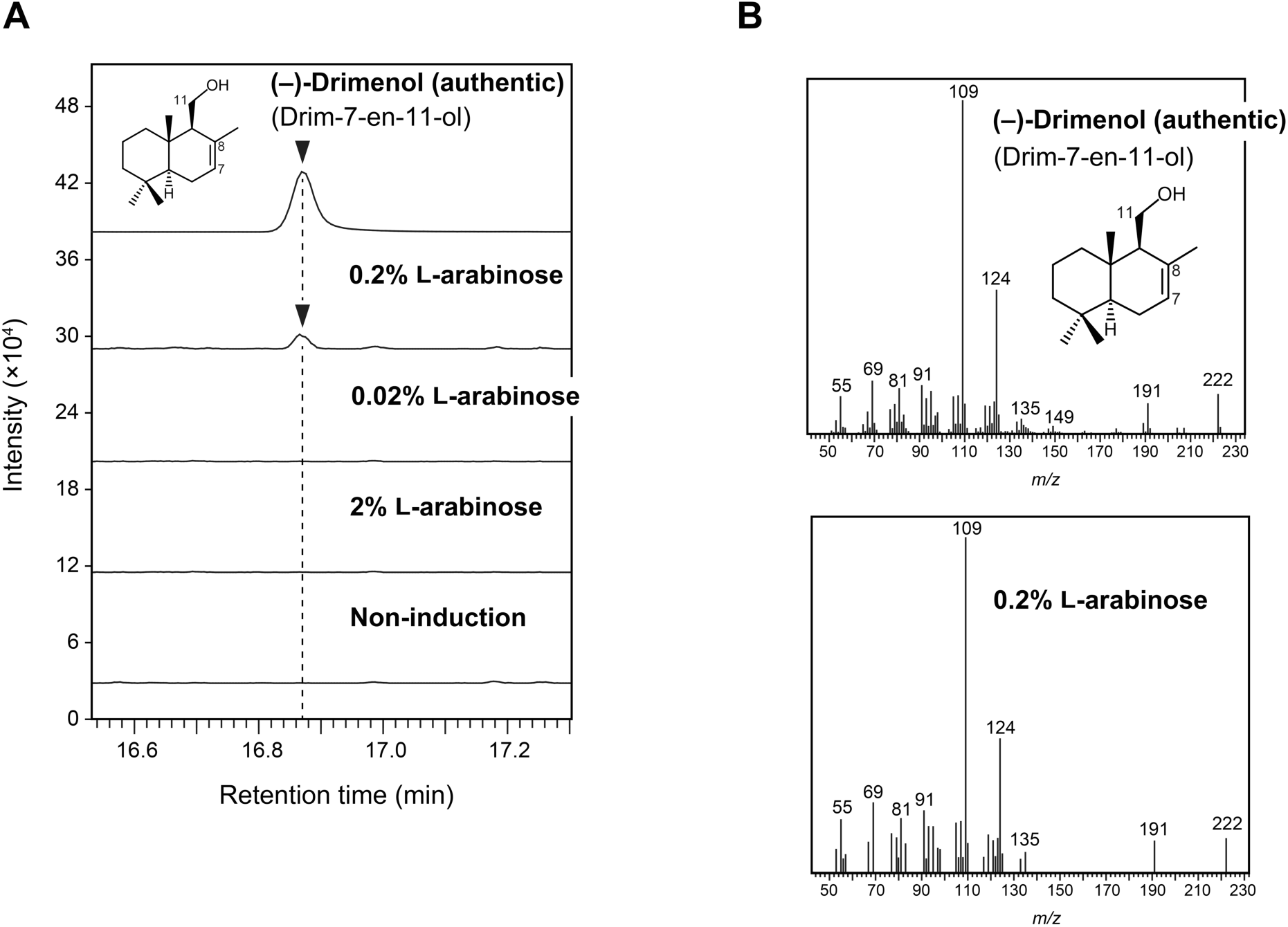
Biosynthesis of (–)-drimenol in *F*. *eckloniae* under L-arabinose-inducing conditions. (A) GC-MS chromatograms of authentic (–)-drimenol (drim-7-en-11-ol) and metabolites extracted from *F*. *eckloniae* cultured in marine broth medium supplemented without (non-induction) or with 0.02%, 0.2%, or 2% L-arabinose. (B) Mass spectra of authentic drimenol and products extracted from *F*. *eckloniae* cultured with 0.2% L-arabinose detected in (A). The GC-MS data in (A) represent three independent experiments. *m/z*, mass-to-charge ratio.

### Distribution of the Drimenol Biosynthetic Gene Cluster Unique to Marine Bacteria

We questioned whether the drimenol biosynthetic gene clusters are either highly conserved or unique within bacterial genomes. To investigate the evolutionary trajectory of the drimenol gene clusters, we searched for any other possible drimenol clusters in bacteria and found that bacteria that harbored the drimenol gene cluster were isolated from a variety of marine habitat samples, collected mainly from surface waters (Figure 4). These data support the idea of terpene biosynthetic gene cluster enrichment in surface or deeper sunlit marine communities in the global open ocean microbiome, described by Paoli *et al*.^6^ The drimenol gene cluster was identified as belonging to family Flavobacteriaceae, and found to remain consistently in the same position; this strong conservation of gene order indicates the early acquisition of these genes during the evolution of Flavobacteriaceae, as well as the existence of a precise regulatory program for the assembly of drimenol cluster genes. However, this is not the case for *Muricauda* sp. SCSIO 64092 genome, which lacks the homologous *DMS* gene. Additionally, we detected high conservation of *AraC*-like regulatory genes in bacteria; these genes are generally involved in the regulation of diverse microorganism biological functions, such as carbon metabolism,^23^ stress responses,^24^ and virulence.^25^

**Figure 4.**
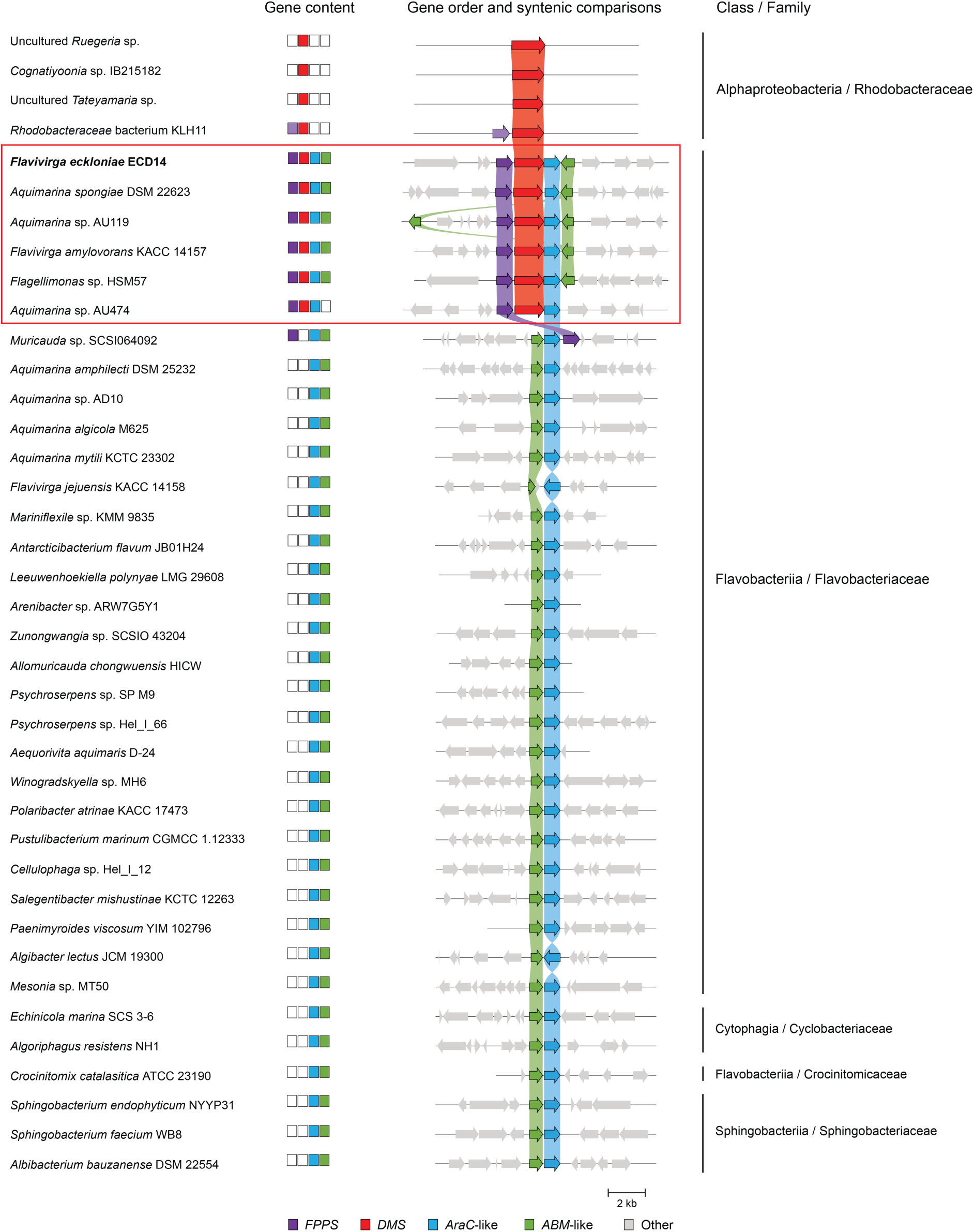
Distribution of drimenol gene clusters across bacteria, with class and family indicated. Central panel shows the presence (colored squares) or absence (white squares) of drimenol cluster genes. Right panel shows the drimenol cluster organization unique to marine bacteria including *Flavivirga eckloniae* (bold black font), and its syntenic comparisons in other bacteria. Red box indicates the conserved drimenol gene cluster in marine bacteria. Genes encoding proteins with similar putative functions with amino acid sequence identity > 30 % are connected by a ribbon. The putative *Rhodobacteraceae* FPPS (sequence identity < 30 %) is shown in light purple. Genes not present in the drimenol cluster are shown in gray. Abbreviations used to describe putative protein-encoding genes: *FPPS*, farnesyl pyrophosphate synthase; *DMS*, drimenol synthase; *AraC*-like, AraC family transcriptional regulator; *ABM*-like, antibiotic biosynthesis monooxygenase.

Interestingly, the aerobic Flavobacteriaceae family was uniquely associated with higher abundances of drimenol cluster genes and thus the biosynthetic potential of drimane sesquiterpenes, which has not been observed in terrestrial bacteria. Metabolomic data has also suggested the uniqueness of some biosynthetic pathways in specific taxa.^26–27^ Symbiotic interactions between marine bacteria and eukaryotic hosts, such as *F*. *eckloniae* and seaweeds^28^ (marine algae) or *A*. *spongiae* and sponges,^29^ have evolutionary and ecological significance. These bacteria respond to environmental changes with diffused specialized metabolites including sesquiterpenes, which act as defensive molecules or a chemical language, mediating communication among bacteria and across domains,^7^ conferring a survival advantage for ecological adaptation.

### Parallel Evolution of the Drimenol Biosynthesis Pathways in Bacterial, Fungal, and Plant Species

Fungal and plant species have been reported to contain clusters of drimane sesquiterpene biosynthesis genes, and to produce drimenol and its derivatives.^11,13–16^ However, DMS enzymes of fungal and bacterial origin are phylogenetically distant from those from plants (Figure 5A and Supplementary Figure S9). Considering the catalytic motifs of DMSs,^30^ enzymes found in bacteria and fungi consist of not only DDxxD/DDxxE motifs (HAD-like domain) but also the DxDTT motif (terpene synthase β domain) typical of class II terpene synthases (Figure 5B), whereas those of plant origin conserve only the DDxxD/EExxD motifs (terpene synthase α domain) characteristic of class I terpene synthases (Figure 5C). The drimenyl pyrophosphate synthase SsDMS, which has low sequence identity (< 20 %) to known bacterial DMSs, was identified within the squalene–hopene synthase superfamily and found to conserve only the DxDDT motif (terpene synthase βγ didomain) characteristic of class II terpene synthases (Figure 5A, B).^30–31^ Interestingly, we found different domain assemblies in the DMSs of marine bacteria and fungi (HAD-β didomain), terrestrial bacteria (βγ didomain), and plants (α domain) (Figure 5D). With respect to drimenol biosynthesis pathways, fungi involved both DMSs and tailoring enzymes such as the P450s and dehydrogenases necessary for modifying drimane scaffolds (Figure 5E), as described in the biosynthesis of astellolides in *Aspergillus oryzae*^13^ or calidoustenes in *Aspergillus calidoustus*.^14^ Conversely, drimenol biosynthesis in marine bacteria lacks those modifying enzymes, but has evolved a cluster of closely co-localized genes including *FPPS*, *DMS*, and an *AraC*-like regulator. These substantial differences in cluster gene organization and the sequence identity of relevant enzymes indicates parallel evolution of the ability to produce drimenol in bacteria to that in fungal and plant species.

**Figure 5.**
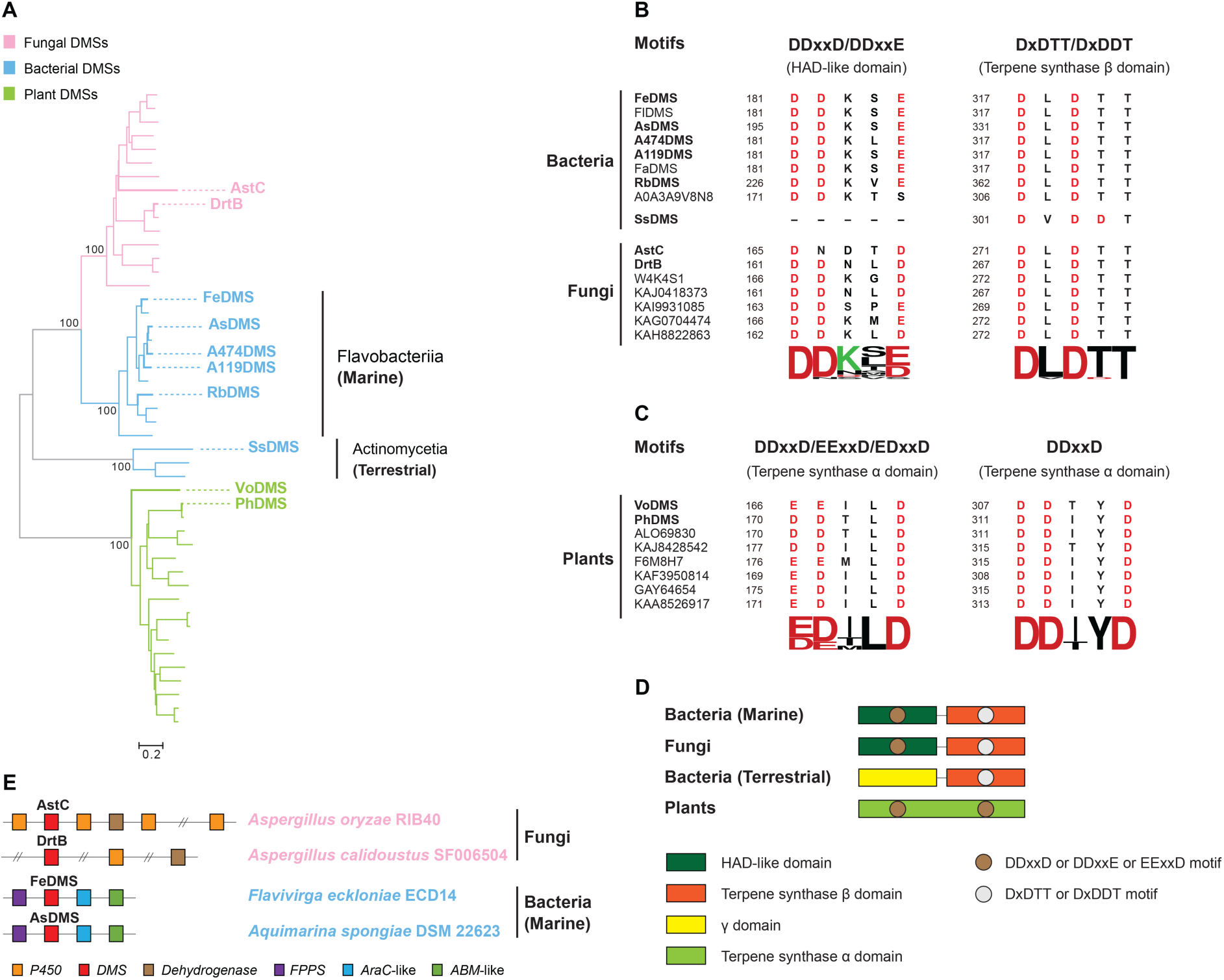
Parallel evolution of the drimenol biosynthesis pathways in bacterial, fungal, and plant species. (A) Neighbor-joining phylogenetic tree of DMS enzymes from bacteria (blue, classes Flavobacteriia and Actinomycetia), fungi (pink), and plants (green). The *bar* represents 0.2 amino acid substitutions per site; *numbers* next to branches are the percentages of replicate trees in which associated taxa clustered together in bootstrap tests with 1,000 replicates. The detailed tree is provided in Supplementary Figure S9. (B–C) Sequence alignments of conserved catalytic motifs of selected DMSs in bacteria and fungi (B) and plants (C). Protein sequences were aligned using GenomeNet ClustalW v1.83 (https://www.genome.jp/tools-bin/clustalw), and sequence logos were generated using the WebLogo server.^32^ Previously reported DMSs^10-^ 14,31 shown in (A) are indicated in bold in (B) and (C). The accession numbers and domains are shown. (D) Domain architectures of DMSs from bacteria (marine and terrestrial), fungi, and plants. Catalytic motifs are indicated. (E) Comparison of gene cluster organization for drimenol in fungi (pink) and marine bacteria (blue). *P450*, cytochrome P450 monooxygenase; *DMS*, drimenol synthase; *FPPS*, farnesyl pyrophosphate synthase; *AraC*-like, AraC family transcriptional regulator; *ABM*, antibiotic biosynthesis monooxygenase. Fe, *Flavivirga eckloniae*; As, *Aquimarina spongiae*; A474, *Aquimarina* sp. AU474; A119, *Aquimarina* sp. AU119; Rb, *Rhodobacteraceae* KLH11; Fl, *Flagellimonas* sp. HSM57; Fa, *Flavivirga amylovorans*; Ss, *Streptomyces showdoensis*; Vo, *Valeriana officinalis*; Ph, *Persicaria hydropiper*.

### Reconstruction of the Evolution of Drimenol Biosynthetic Gene Cluster in Marine Bacteria and Discovery of a Mono-β-domain Terpene Synthase

Our previous study on the characterization of AsDMS showed the required assembly of the two active domains, including a HAD-like domain (N-domain) and terpene synthase β domain (C-domain), for bifunctional catalysis of the AsDMS enzyme.^10^ Therefore, the discovery of monodomain proteins that encode discrete HAD and terpene synthase β may contribute to our understanding of the evolution of the fusion AsDMS enzyme and drimenol biosynthetic gene cluster unique to marine bacteria. Accordingly, we conducted a Basic Local Alignment Search Tool (BLAST) search for bacterial genomes in the NCBI database, using the domain swapping sequence (C– N domain) of the AsDMS protein as a search string. Interestingly, we found individual *HAD* and *terpene synthase β* genes (accession numbers WP_129575347 and WP_129572421, respectively) isolated within the genome of *Sorangium cellulosum* strain Soce836 (Figure 6A), a soil-dwelling bacterium of the Polyangiaceae family. By tracing the phylogenetic relationship of the HAD enzyme of *S*. *cellulosum* (Sc HAD) with the N-domain of AsDMS and known HAD-like hydrolases, we demonstrated that the Sc HAD protein was homologous to the N-domain of AsDMS and clustered with bacterial HAD phosphatases (Figure 6B). Furthermore, an amino acid alignment comparison revealed that Sc HAD conserves four short HAD signature motifs^33–34^ including the DDxxE_168–172_ motif (HAD motif IV) that was also found in AsDMS (DDxxE_195–199_ motif)^10^ and class I terpene synthases (Supplementary Figure S10A). Next, we constructed the phylogeny of the terpene synthase β enzyme from *S*. *cellulosum* (Sc terpene synthase β) with the C-domain of AsDMS and functionally characterized terpene synthases of β domains, and found that Sc terpene synthase β was clustered with bacterial class II terpene synthases and homologous to the C-domain of AsDMS (Figure 6C). Extensive sequence comparisons showed that the DxDVT_134–138_ motif in Sc terpene synthase β corresponded closely to the DxDTT_331–335_ motif (C-domain) of AsDMS^10^ or DxDTT/DxDDT motifs of other enzymes (Figure 6D and Supplementary Figure S10B).

Comparisons of the sequence and structural model of Sc terpene synthase β constructed using the automated SWISS-MODEL pipeline with those of the C-domain_235–536_ of AsDMS^10^ demonstrated conserved active site-defining amino acid residues within the two proteins, including aspartate residues (D134 and D136 of the DxDVT_134–138_ motif in Sc terpene synthase β versus D331 and D333 of the DxDTT_331–335_ motif in AsDMS) and nearby residues with hydrophobic or positively charged side chains (Y176, R188, Y234, and Y235 in Sc terpene synthase β versus Y373, R380, Y426, and Y427 in AsDMS) (Figure 6D, E). The SWISS-MODEL-based structure of Sc HAD also shared high structural similarity with the HAD-like domain (N-domain) of AsDMS^10^ (Figure 6E). These observations indicate the existence of such a mono-β-domain terpene synthase as Sc terpene synthase β, and that both the Sc HAD and Sc terpene synthase β enzymes might be related to the evolution of domain assembly in AsDMS.

Some additional drimenol cluster component genes such as *FPPS*, the *AraC*-like regulator, and *ABM* were present in the *S*. *cellulosum* genome, denoted as *ScFPPS*, *ScAraC*-like, and *ScABM* (Figure 6A); these protein-encoding genes were homologs of corresponding genes from *A*. *spongiae*. Multiple copies of the *FPPS* cluster gene were also present in *ScFPPS1*–*4* in *S*. *cellulosum*, implying redundant or alternative gene functions. A syntenic comparison between the *S*. *cellulosum* and *A*. *spongiae* genomes suggested subsequent gene loss and rearrangement in the drimenol cluster of *A*. *spongiae* (Figure 6A).

**Figure 6.**
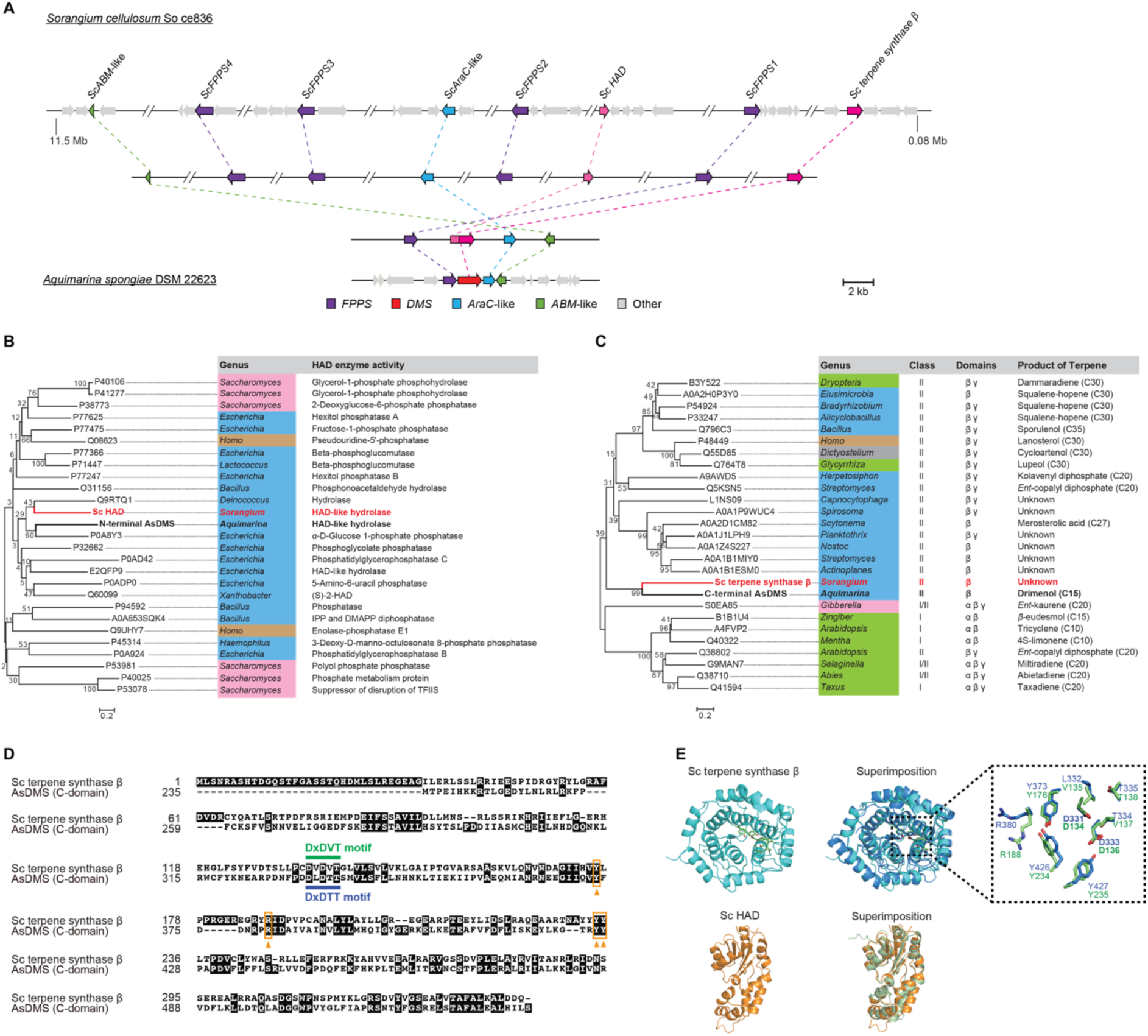
Scenario for the evolution of the drimenol cluster in bacteria. (A) Evolutionary events of the drimenol cluster inferred by syntenic comparison between genomes of *S*. *cellulosum* strain Soce836 and *A*. *spongiae*. Dashed coloured lines indicate putative gene rearrangement events during evolution, and genes encoding for proteins of similar putative functions are noted by the same color. *FPPS*, farnesyl pyrophosphate synthase; *DMS*, drimenol synthase; *AraC*-like, AraC family transcriptional regulator; *ABM*, antibiotic biosynthesis monooxygenase. (B) Neighbor-joining phylogenetic tree of the Sc haloacid dehalogenase (HAD) (bold red text) with the N-terminal domain of AsDMS (bold black text) and functionally characterized HAD-like hydrolases. (C) Neighbor-joining phylogenetic tree of the Sc terpene synthase β (bold red text) with the C-terminal domain of AsDMS (bold black text) and selected β domains from functionally characterized and unknown terpene synthases. The *bar* indicates 0.2 amino acid substitutions per site; *numbers* next to branches are the percentages of replicate trees in which associated taxa clustered together in bootstrap tests with 1,000 replicates. Colors indicate different organisms: bacteria – blue, fungi – pink, plants – green, protists – grey, and animals – light brown. (D) Sequence alignment of the Sc terpene synthase β and C-domain235–536 of AsDMS. The conserved DxDVT and DxDTT motifs are indicated in dark green and dark blue, respectively. The positions of substrate-binding site amino acid residues are shown in the orange boxes with orange *arrowheads*. (E) Upper panel (left to right): an overview map of the SWISS-MODEL-based structure of Sc terpene synthase β (cyan) and its comparison with the AlphaFold2 structure of the C-domain of AsDMS (blue). A close-up of the spatial arrangement of the catalytic DxDVT134–138 motif in Sc terpene synthase β and the DxDTT331–335 motif in AsDMS with substrate-binding site residues is shown in the black-dotted box. The side chains of AsDMS residues (dark blue text) are displayed in atomic coloring (blue, carbon; red, oxygen), whereas those in Sc terpene synthase β (dark green text) are shown in green for carbon and red for oxygen. Catalytically essential aspartate residues are shown in bold black text. Lower panel (left to right): SWISS-MODEL-based structure of the Sc HAD (orange) and its superimposition with the AlphaFold2 structure of N-domain of AsDMS (pale green). Sc, *Sorangium cellulosum*; As, *Aquimarina spongiae*.

### Functional Analysis of the Mono-β-domain Sesquiterpene Synthase

To examine the catalytic function of the individual *S*. *cellulosum* HAD and terpene synthase β enzymes, we expressed these recombinant N-terminal His_8_-tag-fused proteins in *Escherichia coli*, and obtained the purified Sc HAD and terpene synthase β proteins (Figure 7A) for *in vitro* enzymatic activity assays in the presence of FPP as a substrate. The treatment of reaction mixtures with or without BAP was to determine whether the Sc terpene synthase β and Sc HAD proteins were monofunctional in catalyzing FPP cyclization and the dephosphorylation of cyclic diphosphate intermediates, respectively. When Sc terpene synthase β was incubated alone with FPP in the absence of BAP, no cyclization products were detected (Figure 7B). However, the addition of BAP favored dephosphorylation activity, resulting in the formation of a single compound corresponding to a cyclized sesquiterpene alcohol in terms of its retention time (15.688 min) and mass fragmentation pattern (parent mass *m/z* = 222) (Figure 7B, C), confirming that Sc terpene synthase β is monofunctional and performs only the cyclization step of FPP. Furthermore, the Sc terpene synthase β enzyme did not require the addition of Mg^2+^ for its sesquiterpene synthase activity (Supplementary Figure S11), which has not previously been observed in known sesquiterpene synthases.^19,31^ Next, we performed coupled enzymatic activity assays for Sc HAD (0.2–1 *μ*M, Figure 7B) with Sc terpene synthase β using the FPP substrate; however, we did not observe any sesquiterpene alcohol in the absence of BAP treatment (Figure 7B). The lack of dephosphorylation activity of the Sc HAD enzyme may be explained by its large distance from the Sc terpene synthase β enzyme on the *S*. *cellulosum* genome, suggesting that it required evolutionary events to diversify its protein functions.

**Figure 7.**
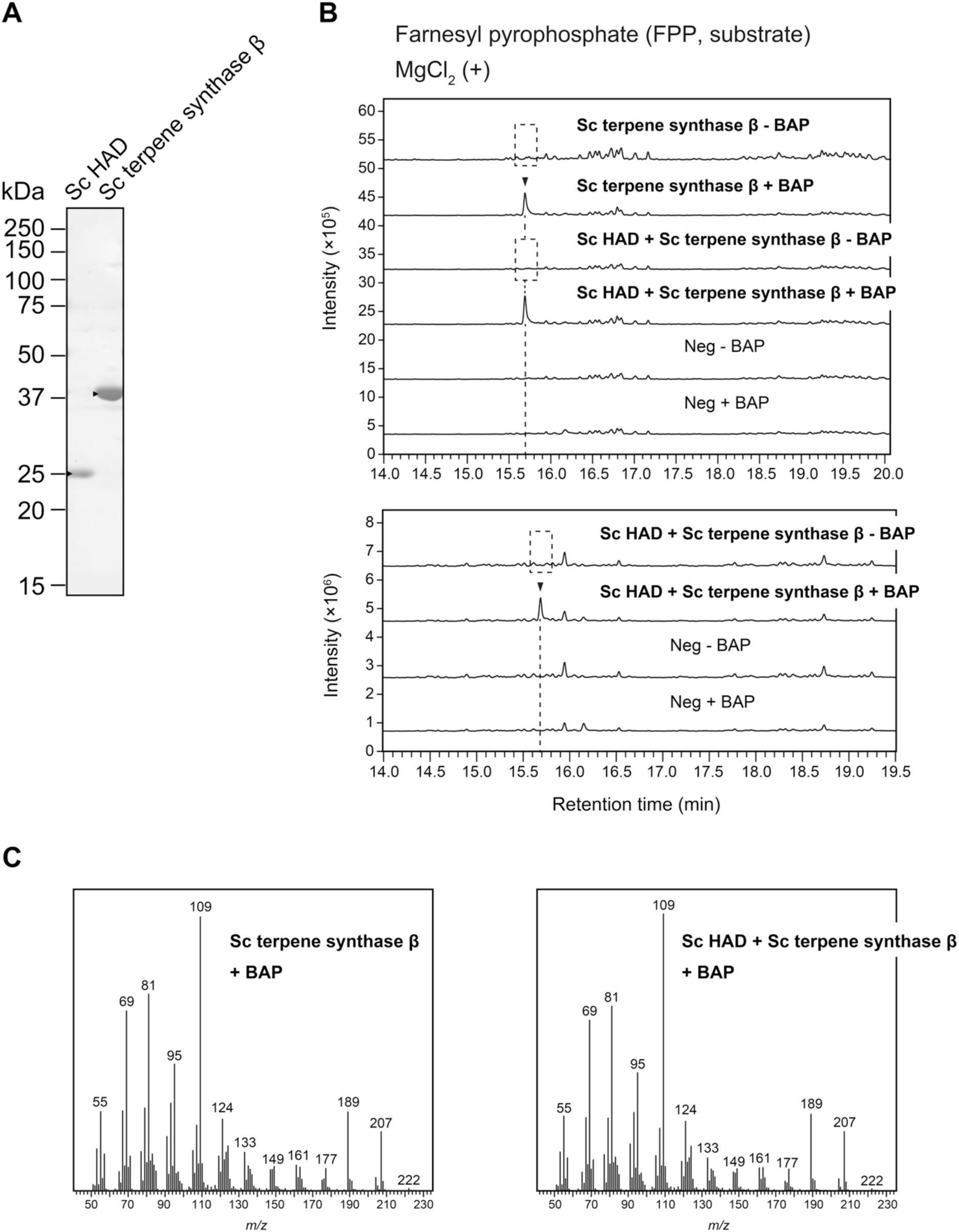
Catalytic activity of the recombinant Sc HAD and Sc terpene synthase β proteins. (A) 12.5% sodium dodecyl sulfate–polyacrylamide (SDS-PAGE) analysis of purified Sc HAD and Sc terpene synthase β proteins (1 *μ*g) from *E*. *coli*. The gel was stained with Coomassie Brilliant Blue, and the positions of molecular markers are indicated. *Arrowheads* indicate the desired protein bands. (B) GC-MS analyses of *in vitro* reaction products. Enzymatic assays of purified Sc terpene synthase β protein alone or coupled with the Sc HAD (0.2 *μ*M, upper panel) and the Sc terpene synthase β protein coupled with Sc HAD (up to 1 *μ*M, lower panel) were performed individually in the presence of farnesyl pyrophosphate (FPP) substrate, with or without alkaline phosphatase (BAP) treatment. Reactions in the absence of Sc terpene synthase β and Sc HAD proteins were used as negative controls (Neg). The GC-MS data represent three independent reactions performed with the same preparation of purified protein. (C) Mass spectra of reaction products detected in (B). Dashed boxes indicate that no desired reaction products were detected. Sc, *Sorangium cellulosum*. *m/z*, mass-to-charge ratio.

To our knowledge, this is the first study to demonstrate monofunctional cyclization activity of a single-β-domain class II terpene synthase in the biosynthesis of sesquiterpenes, an architecture previously unseen for sesquiterpene synthases, which are traditionally known to contain βγ or αβγ.^19,35–36^ Our findings reinforce the structural and functional divergence of natural terpene synthases. Further studies on the rational engineering of monodomain proteins via acquisition of new functional domains derived from different enzyme origins to create new catalytic centers and diversify protein functions are awaited. Indeed, the *A*. *spongiae* AsDMS enzyme was recently observed to fuse the HAD-like domain with the terpene synthase β domain and displays its bifunctional activity.^10^

## CONCLUSIONS

In this study, we elucidated the genetically encoded drimenol biosynthetic pathway in marine bacteria upon arabinose induction, demonstrated the extent of drimenol biosynthetic potential and its genomic context across bacteria, and discovered the unique gene arrangement and composition of the drimenol cluster in marine bacteria, which may have evolved in parallel to those of plants and fungi. Notably, our findings of the monodomain HAD and terpene synthase β proteins of *S*. *cellulosum* suggest domain fusion during the evolution of drimenol biosynthesis in bacteria. These findings highlight the unique biosynthetic potential of drimane sesquiterpenes in marine bacteria and demonstrate the evolutionary trajectory of unusual terpene synthase architectures, which will inspire future efforts to rationally engineer new catalytic functions via domain fusion of bacterial enzymes.

## EXPERIMENTAL SECTION

### Genomic Data Mining for Drimenol Biosynthetic Genes in *F*. *eckloniae* and Other Marine Bacteria

Because no drimenol biosynthetic gene clusters in bacteria were predicted using the AntiSMASH software,^37^ we investigated the gene neighborhoods of *FeDMS* for the clustered genes, using the genome sequence of *F*. *eckloniae* in the NCBI database (accession number CP025791). The putative proteins encoded in the cluster were characterized using the PFAM and InterPro protein family databases. The sequences of 10 upstream and 10 downstream genes adjacent to the drimenol cluster were also extracted and inspected for protein families. Similarly, drimenol biosynthetic genes in different marine bacteria containing the *DMS* gene were identified against their genome sequences, including *A*. *spongiae* (FQYP01000002), *Aquimarina* sp. AU119 (OMPF01000021), *F*. *amylovorans* (JAUOEM010000005), *Flagellimonas sp. HSM57* (WHOT01000003), *Aquimarina* sp. AU474 (OMPE01000007), and *Rhodobacteraceae* KLH11 (DS999533). The representations of all putative drimenol gene clusters were investigated using Integrative Genomics Viewer v2.8.2,^38^ and diagrams were created using CAGECAT v1.0.^39^ The protein sequences of FeFPPS, AsFPPS, FeABM, AsABM, FeAraC, and AsAraC are available in the NCBI database (accession numbers WP_102757878, WP_073315021, WP_102757881, SHI65374, WP_102757880, and WP_170864563, respectively).

### Materials for Cloning and Enzyme Activity Assays

The bacterial strains, plasmids, chemicals, and enzymes are described in Method S2.

### Plasmid Construction, Expression, and Purification of FPPS, Sc HAD, and Sc terpene synthase β Proteins

Coding sequences of candidate *FPPS*, *Sc HAD*, and *Sc terpene synthase β* genes were codon-optimized for expression in *E*. *coli* and synthesized by Eurofins Genomics (Tokyo, Japan). Each sequence was ligated into the *Nde* I and *EcoR* I sites of the linearized pET28b(+) expression vector containing N-terminal octa-histidine-tag (His_8_-tag), followed by transformation into *E*. *coli* BL21 Star (DE3). The empty pET28b(+)-His_8_ vector was used as a negative control for protein expression.

All recombinant N-terminal His_8_-tagged FPPS proteins were expressed in *E*. *coli* BL21 Star (DE3) strains harboring pET28b(+)-His_8_*FPPS* and purified using TALON metal affinity resin (Clontech, Mountain View, CA, USA), and their concentrations were determined, as previously described.^10^ Expression and purification of the Sc HAD and Sc terpene synthase β proteins were conducted as described for the recombinant FPPS proteins. These purified recombinant proteins were examined using sodium dodecyl sulfate polyacrylamide (SDS-PAGE) and used for *in vitro* enzymatic activity assays.

### *In Vitro* Enzymatic Activity Assays

For the characterization of the FPPS proteins, the reaction was initiated by the addition of purified proteins (FeFPPS or AsFPPS, final 200 nM) to a mixture of 50 mM Tris-HCl (pH 8.0), 2 mM MgCl_2_, 1 mM dithiothreitol, 100 *μ*M IPP, and 100 *μ*M DMAPP or GPP. Reactions in the absence of proteins were used as negative controls. The reaction mixture (200 *μ*L) was incubated at 30 °C for 1 h, followed by the addition of BAP from *E*. *coli* C75 (4 *µ*L, 2 units; Takara Bio) and further incubation for 1 h at 37 °C. For the coupled enzymatic activity assay of FPPS and DMS proteins, a reaction mixture with a total volume of 200 *μ*L, containing 50 mM Tris-HCl (pH 8.0), 4 mM MgCl_2_, 2 mM dithiothreitol, 100 *μ*M IPP, 100 *μ*M DMAPP or GPP, and purified proteins (FeFPPS or AsFPPS, final 1 *μ*M; FeDMS or AsDMS, final 200 nM) was incubated at 30 °C for 1 h. All reaction products were then extracted twice using 200 *μ*L of hexane/ethyl acetate solution (1:1, v/v) and centrifuged at 4,400 × *g* for 5 min at 4 °C. The resulting upper organic layer was combined and subjected to GC-MS analysis.

The Sc HAD and Sc terpene synthase β proteins were characterized using a mixture (400 *μ*L) containing 50 mM Tris-HCl (pH 8.0), 2 mM MgCl_2_, 1 mM dithiothreitol, 100 *μ*M FPP, and 0.2 *μ*M purified Sc terpene synthase β alone or coupled with 0.2–1 *μ*M purified Sc HAD; the mixture was reacted at 30 °C within 1 h. The Sc terpene synthase β protein was also characterized in the absence of Mg^2+^, as no MgCl_2_ was added to the mixture. The addition of BAP from *E*. *coli* C75 to 200 *μ*L of reaction mixture and the extraction of products for GC-MS analysis were conducted as described for the characterization of FPPS proteins.

### GC-MS Analysis

The detailed analysis and detection of enzyme reaction products was performed using GC-MS with a 7890 GC (Agilent Technologies, Santa Clara, CA, USA) coupled with a 5975 mass selective detector, based on our previously reported method,^10^ but using a modified HP-5ms UI capillary column (30 m × 250 *μ*m × 0.25 *μ*m). The potential reaction products were compared with authentic standards. The amount of product was measured as the peak area using MSD ChemStation G1701EA E.02.02.1431 GC/MS software (Agilent Technologies). Results obtained from three independent experiments were expressed as means ± standard deviation (SD). Standard curves for authentic farnesol and (–)-drimenol were obtained by GC-MS to calibrate the raw data.

### Cultivation of *F*. *eckloniae* and Analysis of Metabolite Content by GC-MS

The *F*. *eckloniae* ECD14T strain (culture JCM 31797) used in this study was purchased from RIKEN BioResource Research Center (Tsukuba, Ibaraki, Japan) and recovered on marine agar plates. The cells were inoculated with 3 mL of marine broth medium and allowed to grow overnight at 25 °C with shaking at 200 rpm. We added 1 mL of this starter culture to 100 mL of marine broth medium overlaid with dodecane (10% of the total culture volume) and incubated for 2 days at 25 °C with shaking at 160 rpm. A 50-*μ*L aliquot of (+)-nootkatone (final concentration, 2 mM) was added as an internal standard. To test the quality of the drimenol extract, a positive control supplemented with drimenol (final concentration, 100 *μ*M) was prepared. Sesquiterpene extraction was performed as previously described.^40^ Briefly, 100 mL of culture broth was mixed with an equivalent volume of acetone. After sonication for 1 min, the acetone broth was extracted with 70 mL of hexane three times, and the hexane extract was concentrated under a gentle nitrogen stream. Next, 2 mL of extract was loaded onto a 1-mL silica gel column and washed with hexane (1 mL) three times. The metabolites were eluted with 2 mL of hexane/ethyl acetate (7:3, v/v) and further analyzed using GC-MS.

To achieve inducing conditions, following the addition of the starter culture to 100 mL of marine broth medium, the cells were allowed to grow for 2 h at 25 °C with shaking at 160 rpm until the optical density at 600 nm (OD_600_) reached ∼0.2. Then, L-arabinose was added to the culture at 0.02%, 0.2%, or 2% to induce *FeDMS* expression, and overlaid with dodecane (10% of the total culture volume). The contents were incubated for 2 days at 25 °C with shaking at 160 rpm. A non-inducing culture without arabinose was used as a control sample. Metabolite extraction and GC-MS analysis were performed as described above. All assays were conducted in triplicate.

### Homology Searches and Distribution of the Drimenol Cluster in Bacterial Genomes

To investigate bacterial drimenol clusters, we conducted cblaster^41^ analysis against bacterial genomes representing bacterial diversity across terrestrial, deep ocean, and surface seawater habitats in the NCBI database, using the drimenol gene cluster in *F*. *eckloniae* or *A*. *spongiae* as a query. Cluster diagrams were created using the clinker online tool.^42^ Drimenol clusters were detected in bacterial isolates based on the presence of at least three co-occurring homologous genes with a sequence identity of > 30% to those from the drimenol cluster of *F*. *eckloniae* or *A*. *spongiae*.

### Inference of the Evolution of Drimenol Cluster

To infer the evolution of the drimenol cluster across bacteria, we used the tBLASTn program with the domain swapping sequence (C–N domain) of AsDMS as the search string. Evolutionary events affecting the drimenol cluster were analyzed through syntenic comparison between the *S*. *cellulosum* and *A*. *spongiae* genomes using the CAGECAT v1.0^39^ web server.

### Phylogenetic Analyses

Single-gene trees were generated by querying each encoded protein of the drimenol cluster in the NCBI database. These trees were aligned using the GenomeNet ClustalW v1.83 server (https://www.genome.jp/tools-bin/clustalw), and single-gene trees were generated with MEGA 7.0,^43^ based on a neighbor-joining method using the *p*-distance algorithm with 1,000 bootstrap replicates. Multiple sequence alignments were colored using pyBoxshade (https://github.com/mdbaron42/pyBoxshade). Sequence logos were generated using the WebLogo server.^32^

### Structural Model Studies

The three-dimensional structures of individual HAD and terpene synthase β proteins from *S*. *cellulosum* Soce836 were predicted by the SWISS-MODEL^44^ pipeline, using AlphaFold DB (https://alphafold.com) models of the putative HAD (A0A150RXN1_SORCE) and the putative prenylcyclase (A0A150QQP3_SORCE) from *S*. *cellulosum* as template structures, respectively. The structures of both N- and C-domains of *A. spongiae* AsDMS were predicted by AlphaFold2.^45^ All structural models were visualized using PyMOL v1.8 (Schrödinger, New York, NY, USA).

## ASSOCIATED CONTENT

### Supporting Information

Metabolite analysis of *F*. *eckloniae* cell extracts under non-inducing conditions; phylogenetic tree and sequence comparison of FeFPPS and AsFPPS with FPPS proteins; phylogeny of FeABM and AsABM with ABM proteins of bacterial and fungal origin; sequence alignment and phylogenetic tree of FeAraC and AsAraC with AraC transcriptional regulators of bacterial origin; expression and catalytic activity of recombinant FeFPPS and AsFPPS proteins; expression of recombinant FeABM and AsABM proteins in *E*. *coli* and *in vitro* enzymatic assays of these proteins; phylogenetic tree of DMS enzymes from bacteria, fungi, and plants; sequence comparisons of HAD and terpene synthase proteins in *S*. *cellulosum* with related proteins; *in vitro* enzymatic assays of Sc terpene synthase β protein in the absence of Mg^2+^; Methods S1 and S2.

## AUTHOR INFORMATION

### Corresponding Authors

Nhu Ngoc Quynh Vo

*Tel: +81-45-503-9111;

Shunji Takahashi

*Tel: +81-48-462-1111;

### Author Contributions

NNQV and ST conceived and designed the research; NNQV and YN performed the experiments; NNQV, YN, and ST analyzed and interpreted the data; and NNQV wrote the manuscript.

### Notes

The authors declare no competing financial interest.

## ACKNOWLEDGEMENTS

This research was supported by JSPS KAKENHI Grant-in-Aid for Early-Career Scientists, grant number 23K13898, and RIKEN Incentive Research Project to NNQV; and by JSPS KAKENHI Grant-in-Aid for Scientific Research(A), grant number 20H00416, and Transformative Research Area(A), grant number 23H04564 to ST.

