## Supporting Information for "Parallel Evolution of a Drimenol Biosynthetic Gene Cluster Unique to Marine Bacteria"

**Table of contents**


| | **Page** |
|---|---|
| **Figure S1.** Metabolite analysis of *F. eckloniae* cell extracts under non-inducing condition. | S2 |
| **Figure S2.** Neighbor-joining phylogenetic tree of FeFPPS and AsFPPS (bold black font) with functionally characterized and unknown FPPSs. | S3 |
| **Figure S3.** Sequence comparison of FeFPPS and AsFPPS (bold black font) with selected FPPS proteins. | S4 |
| **Figure S4.** Neighbor-joining phylogenetic tree of FeABM and AsABM proteins with functionally characterized and unknown ABMs of bacterial and fungal origin. | S5 |
| **Figure S5.** Multiple sequence alignment of *F. eckloniae* FeAraC and *A. spongiae* AsAraC proteins with selected AraC transcriptional regulators of bacterial origin. | S6 |
| **Figure S6.** Neighbor-joining phylogenetic tree of FeAraC and AsAraC proteins with functionally characterized and unknown bacterial AraC-type transcriptional regulators. | S7 |
| **Figure S7.** Expression and catalytic activity of recombinant FeFPPS and AsFPPS proteins. | S8 |
| **Figure S8.** Expression of recombinant FeABM and AsABM proteins in *E. coli* and *in vitro* enzymatic assays of these proteins. | S9 |
| **Figure S9.** Neighbor-joining phylogenetic tree of DMS enzymes from bacteria, fungi, and plants. | S10 |

| | |
|---|---|
| **Figure S10.** Sequence comparisons of HAD and terpene synthase proteins in *S. cellulosum* with selected HAD-like hydrolases and β domains of terpene synthases, respectively. | S11 |
| **Figure S11.** *In vitro* enzymatic assays of Sc terpene synthase β protein in the absence of $Mg^{2+}$. | S12 |
| **Methods S1 and S2** | S13 |


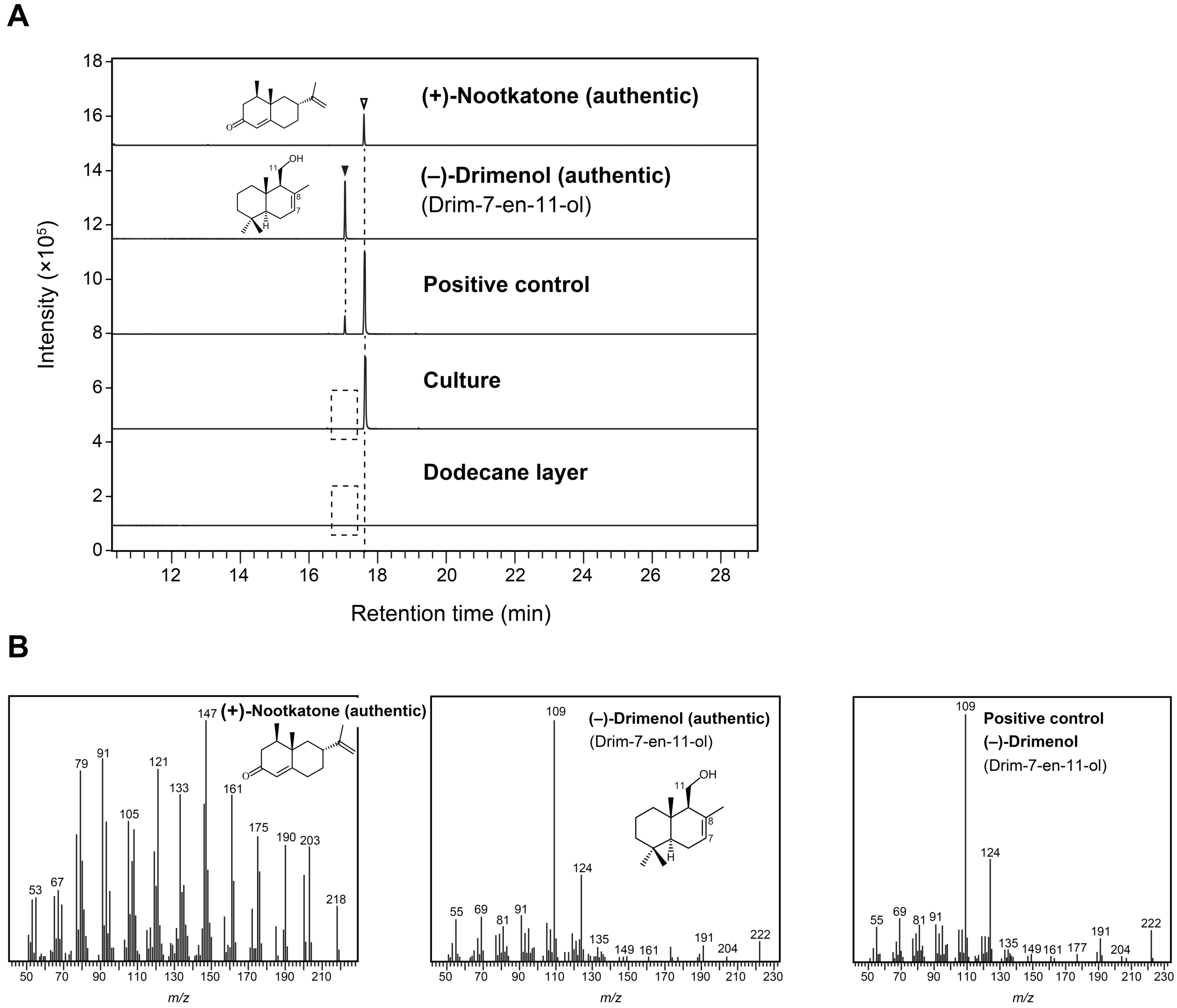


**Figure S1.** Metabolite analysis of *F. eckloniae* cell extracts under non-inducing condition. (A) GC-MS chromatograms of authentic (+)-nootkatone and (–)-drimenol (drim-7-en-11-ol) and metabolites (culture and dodecane layer) from *F. eckloniae* cultured in marine broth medium. The cell extract to which drimenol was added to test the extraction quality was used as a positive control. GC-MS data in (A) represent three independent experiments. Dashed boxes indicate that no desired products were detected. (B) Mass spectra of authentic (+)-nootkatone and (–)-drimenol. *m/z*, mass-to-charge ratio.

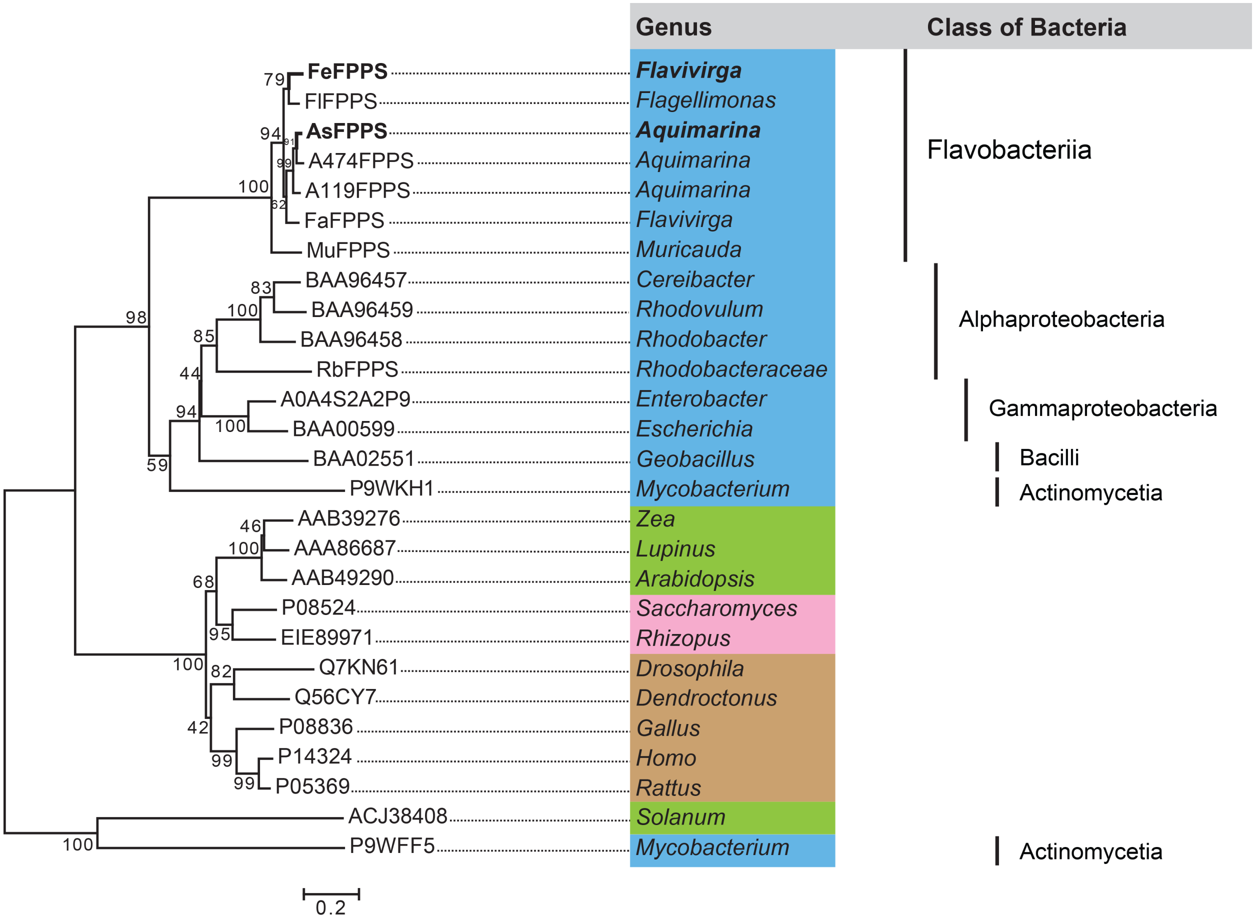


**Figure S2.** Neighbor-joining phylogenetic tree of FeFPPS and AsFPPS (bold black font) with functionally characterized and unknown FPPSs. The bar indicates 0.2 amino acid substitutions per site. The *numbers* shown next to branches are the percentages of replicate trees in which associated taxa clustered together in bootstrap tests with 1,000 replicates. Organisms are colored based on their phylogenetic origin: bacteria – blue, fungi – pink, plants – green, and animals – light brown. Accession numbers and classes of bacteria are indicated. Protein sequences of FlFPPS, A474FPPS, A119FPPS, FaFPPS, MuFPPS, and RbFPPS are available under NCBI accession numbers WP_163419724, WP_109300390, WP_109437249, WP_303283566, WP_247132896, and WP_008759291, respectively. A119, *Aquimarina* sp. AU119; A474, *Aquimarina* sp. AU474; As, *Aquimarina spongiae*; Fa, *Flavivirga amylovorans*; Fe, *Flavivirga eckloniae*; Fl, *Flagellimonas* sp. HSM57; Mu, *Muricauda* sp. SCSIO 64092; Rb, *Rhodobacteraceae* KLH11.

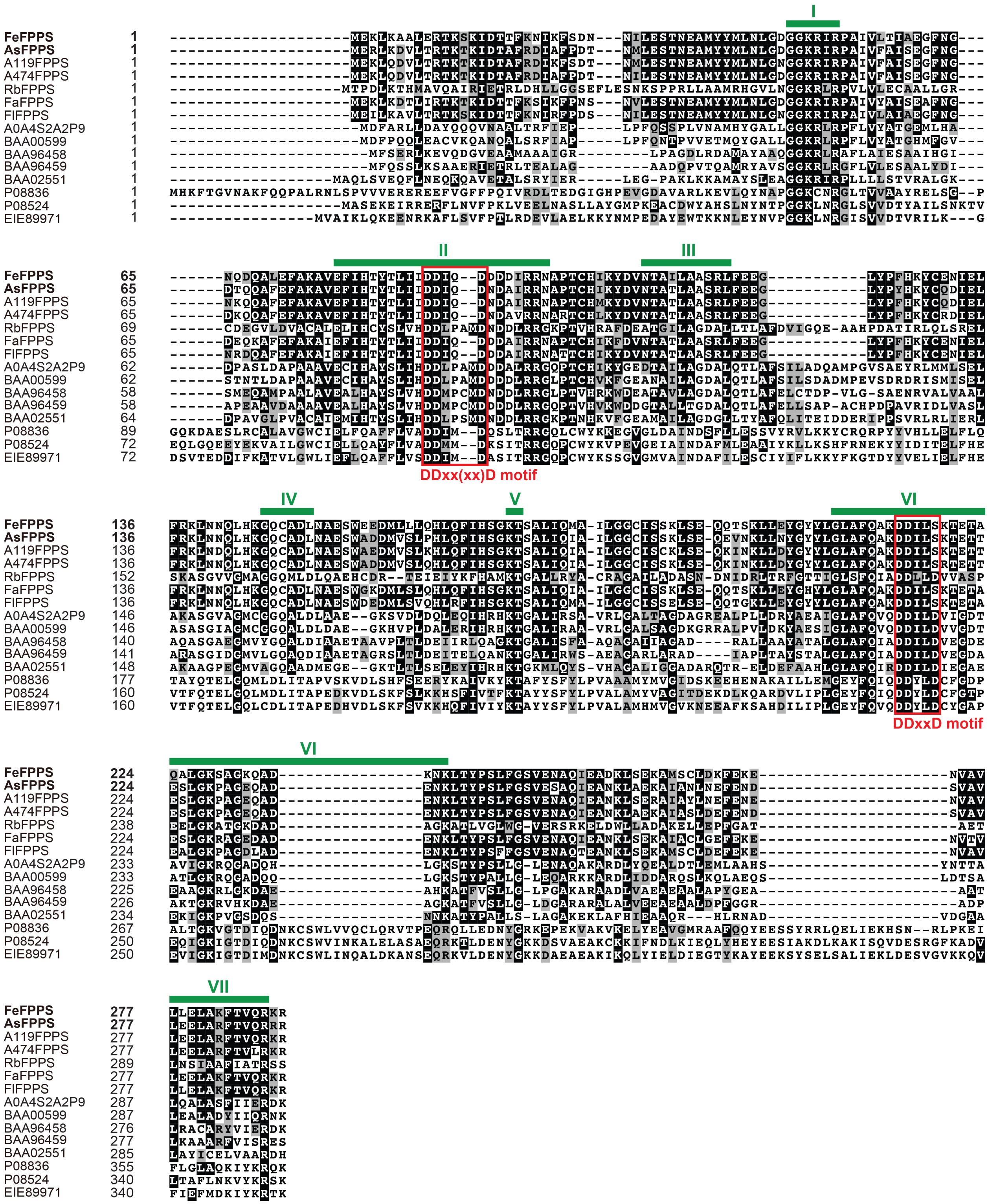


**Figure S3.** Sequence comparison of FeFPPS and AsFPPS (bold black font) with selected FPPS proteins, showing highly conserved regions (I–VII, green bars) typical of FPPSs. The first aspartate-rich motif DDxx(xx)D in region II and the second aspartate-rich motif DDxxD in VI are highlighted in red boxes. Protein sequences were aligned and colored using GenomeNet ClustalW v1.83 (https://www.genome.jp/tools-bin/clustalw) and the BoxShade 3.21 server (https://embnet.vital-it.ch/software/BOX_form.html).

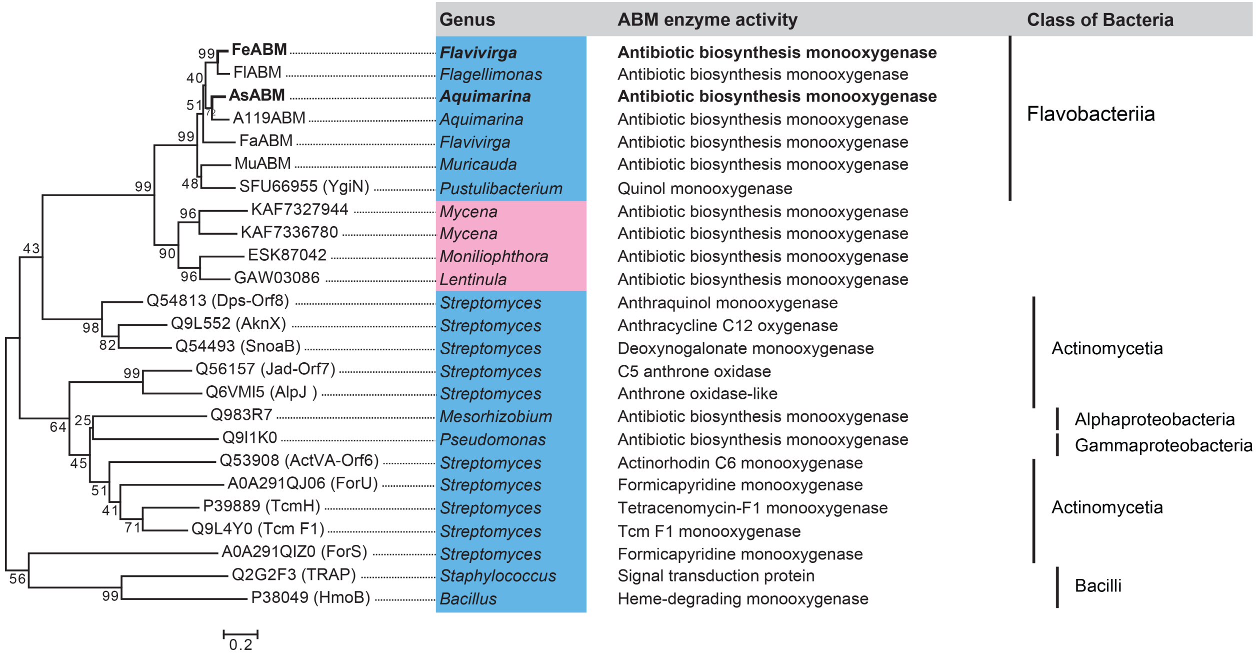


**Figure S4.** Neighbor-joining phylogenetic tree of FeABM and AsABM proteins (bold) with functionally characterized and unknown ABMs of bacterial (blue) and fungal (pink) origin. The bar indicates 0.2 amino acid substitutions per site; *numbers* next to branches are the percentages of replicate trees in which associated taxa clustered together in bootstrap tests with 1,000 replicates. Accession numbers, ABM enzyme activity, and bacterial classes are shown. The protein sequences of FlABM, A119ABM, FaABM, and MuABM are available under NCBI accession numbers WP_163419721, WP_160114244, WP_303283563, and WP_247132894, respectively. As, *Aquimarina spongiae*; A119, *Aquimarina* sp. AU119; Fa, *Flavivirga amylovorans*; Fe, *Flavivirga eckloniae*; Fl, *Flagellimonas* sp. HSM57; Mu, *Muricauda* sp. SCSIO 64092.

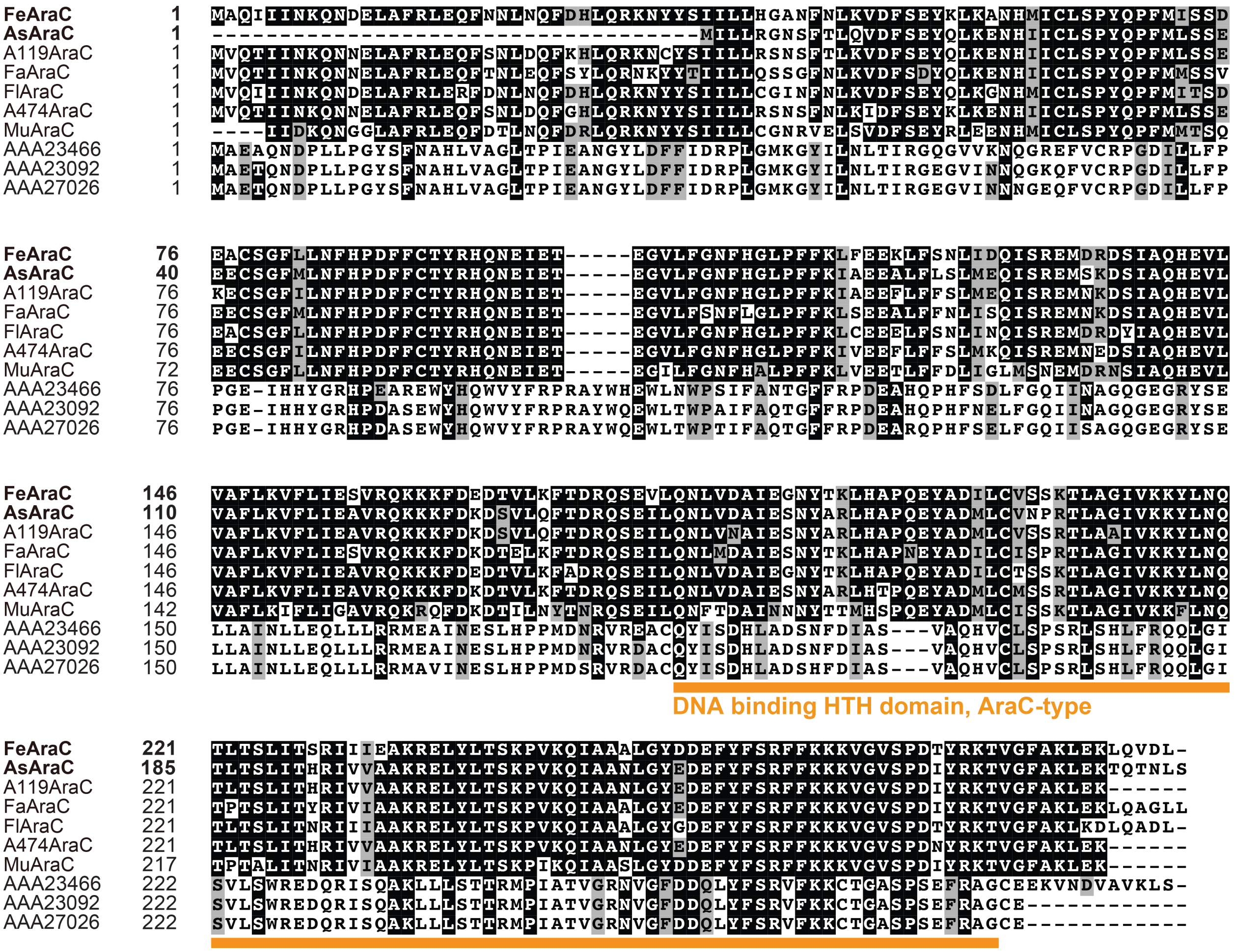


**Figure S5.** Multiple sequence alignment of *F. ecklոniae* FeAraC and *A. spongiae* AsAraC proteins (bold) with selected AraC transcriptional regulators of bacterial origin, showing highly conserved amino acid residues and DNA binding helix-turn-helix (HTH) domain (orange underlining). Sequences were aligned and colored using GenomeNet ClustalW v1.83 and the BoxShade 3.21 server, respectively. Accession numbers are indicated. The NCBI accession numbers for the FlAraC, A119AraC, A474AraC, FaAraC, and MuAraC proteins are WP_163419722, WP_109437247, WP_109300388, WP_303283564, and WP_247132895, respectively. A119, *Aquimarina* sp. AU119; A474, *Aquimarina* sp. AU474; As, *Aquimarina spongiae*; Fa, *Flavivirga amylovorans*; Fe, *Flavivirga ecklոniae*; Fl, *Flagellimonas* sp. HSM57; Mu, *Muricauda* sp. SCSIO 64092.

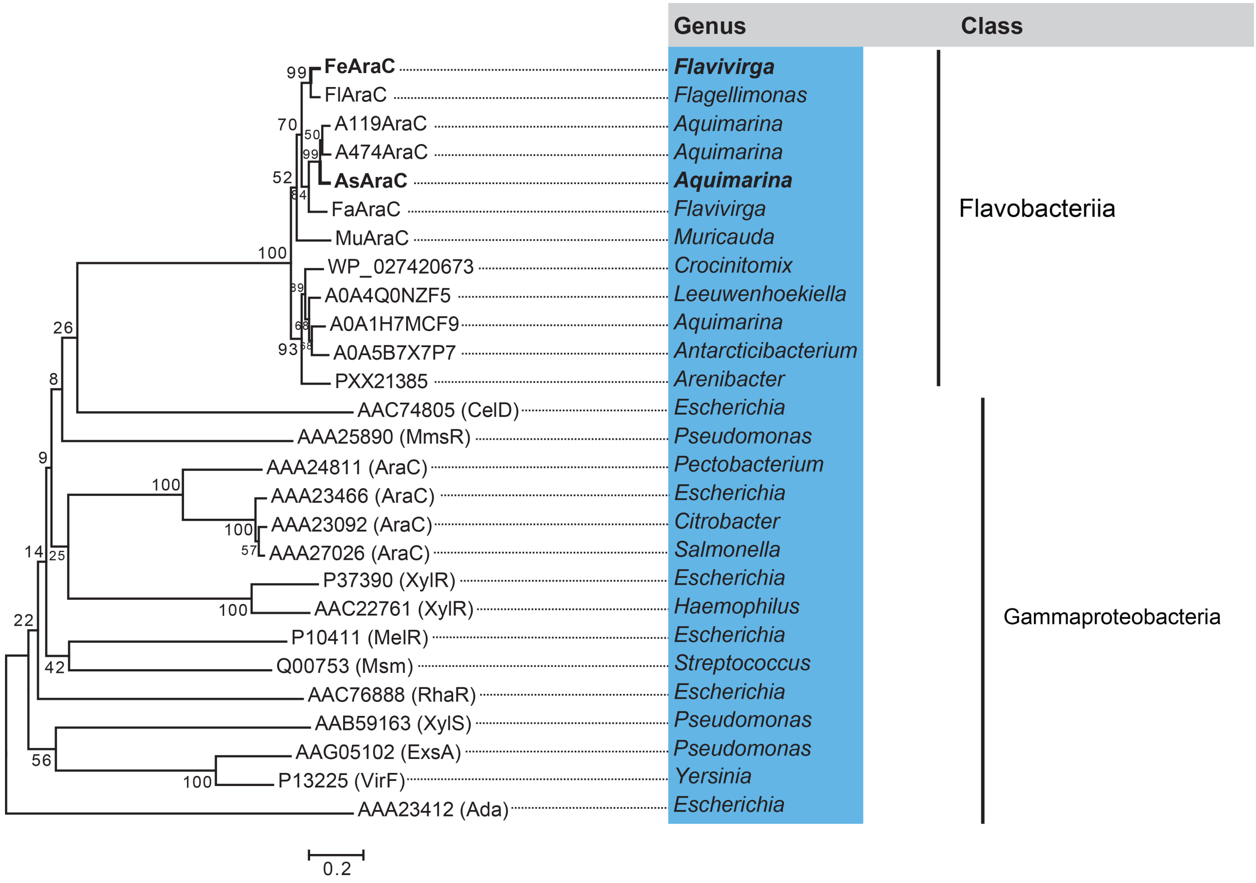


**Figure S6.** Neighbor-joining phylogenetic tree of FeAraC and AsAraC proteins with functionally characterized and unknown bacterial AraC-type transcriptional regulators. The bar indicates 0.2 amino acid substitutions per site; *numbers* next to branches are the percentages of replicate trees in which associated taxa clustered together in bootstrap tests with 1,000 replicates. Bacteria are indicated in blue; their accession numbers and classes are indicated.

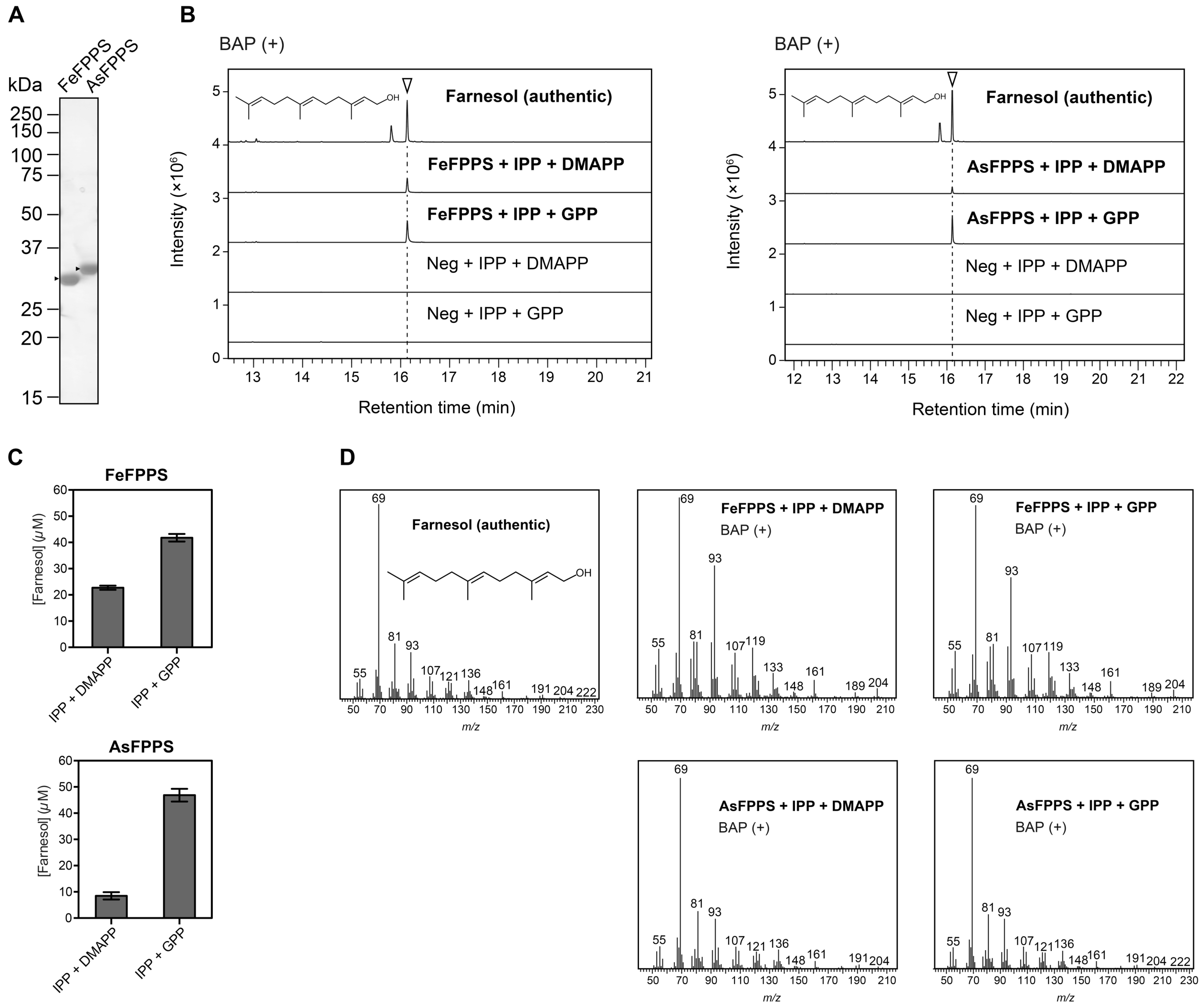


**Figure S7.** Expression and catalytic activity of recombinant FeFPPS and AsFPPS proteins. (A) 12.5% SDS-PAGE analysis of purified FeFPPS and AsFPPS proteins (1 $\mu$g) from *E. coli*. The gel was stained with Coomassie Brilliant Blue, and the positions of molecular markers are indicated. *Arrowheads* indicate FPPS protein bands. (B) GC-MS analyses of authentic farnesol and *in vitro* reaction products. Enzymatic assays of purified FeFPPS (left panel) and AsFPPS (right panel) proteins were performed individually in the presence of substrates: isopentenyl pyrophosphate (IPP) and dimethylallyl pyrophosphate (DMAPP) or geranyl pyrophosphate (GPP), with alkaline phosphatase (BAP) treatment. (C) Comparison of farnesol production after BAP treatment by FeFPPS and AsFPPS proteins. (D) Mass spectra of authentic farnesol and reaction products detected in (B). Data in (B) represent three independent reactions performed with the same preparation of purified protein. Reactions in the absence of FPPS proteins were used as negative controls (Neg). Data in (C) are means ± SD of three independent experiments. As, *Aquimarina spongiae*; Fe, *Flavivirga eckloniae*. *m/z*, mass-to-charge ratio.

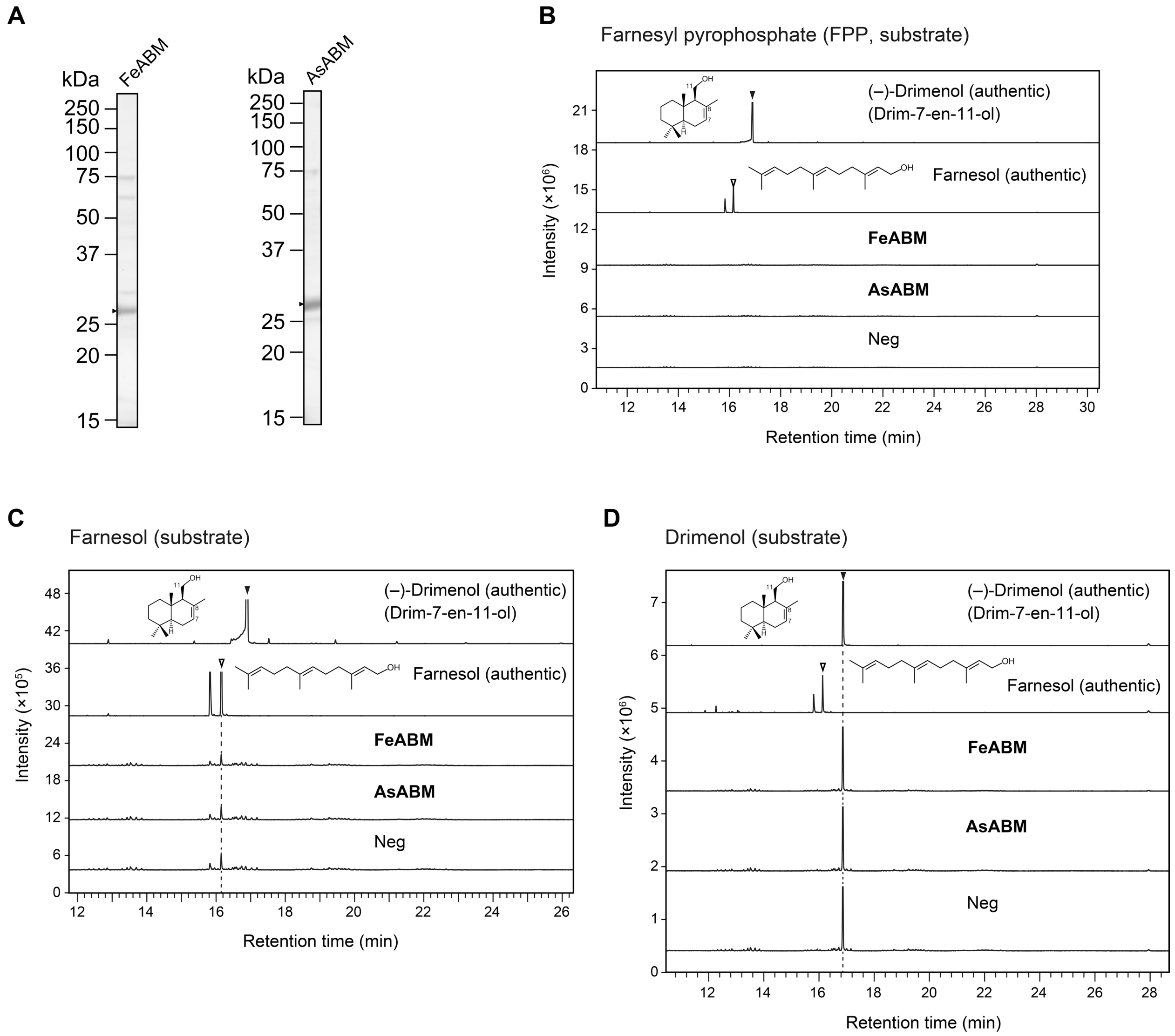


**Figure S8.** Expression of recombinant FeABM and AsABM proteins in *E. coli* and *in vitro* enzymatic assays of these proteins. (A) 12.5% SDS-PAGE analysis of purified FeABM and AsABM proteins (1 $\mu$g) from *E. coli*. The gels were stained with Coomassie Brilliant Blue, and the positions of molecular markers are indicated. *Arrowheads* indicate ABM bands. (B–D) GC-MS chromatograms of authentic (–)-drimenol (drim-7-en-11-ol) and farnesol, and *in vitro* reaction products are shown. Reactions were performed individually in the presence of substrates FPP (B), farnesol (C), or drimenol (D). Reactions in the absence of protein were negative controls (Neg). Data in (B–D) represent three independent reactions performed with the same preparation of purified protein. As, *Aquimarina spongiae*; Fe, *Flavivirga eckloniae*.

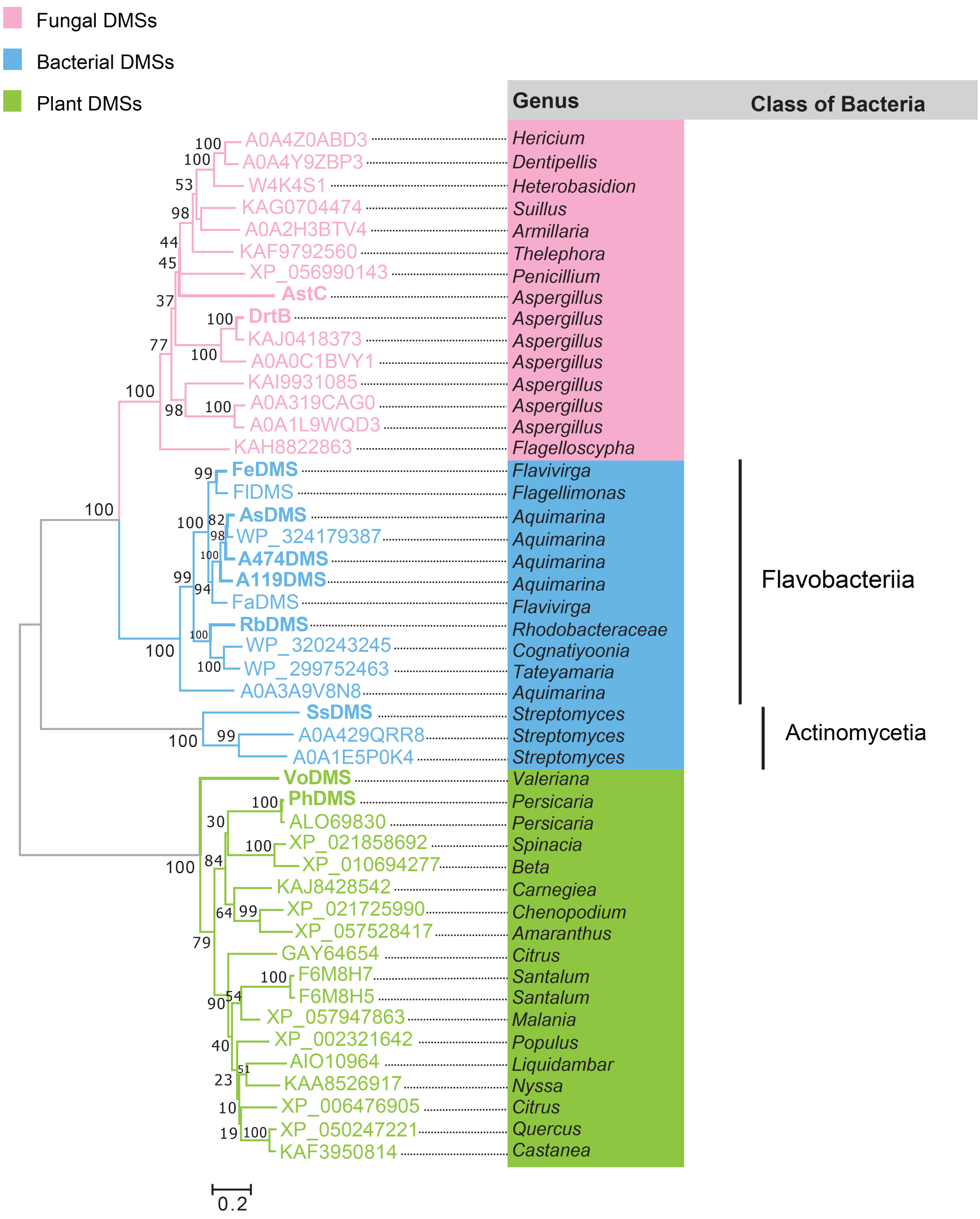


**Figure S9.** Neighbor-joining phylogenetic tree of DMS enzymes from bacteria (blue), fungi (pink), and plants (green). The bar represents 0.2 amino acid substitutions per site; *numbers* next to branches are the percentages of replicate trees in which associated taxa clustered together in bootstrap tests with 1,000 replicates. Accession numbers and classes of bacteria are shown. DMS proteins highlighted in bold are described in Figure 5. Sequences of FlDMS (Fl, *Flagellimonas* sp. HSM57) and FaDMS (Fa, *Flavivirga amylovorans*) are available under NCBI accession numbers WP_163419723 and WP_303283565, respectively.

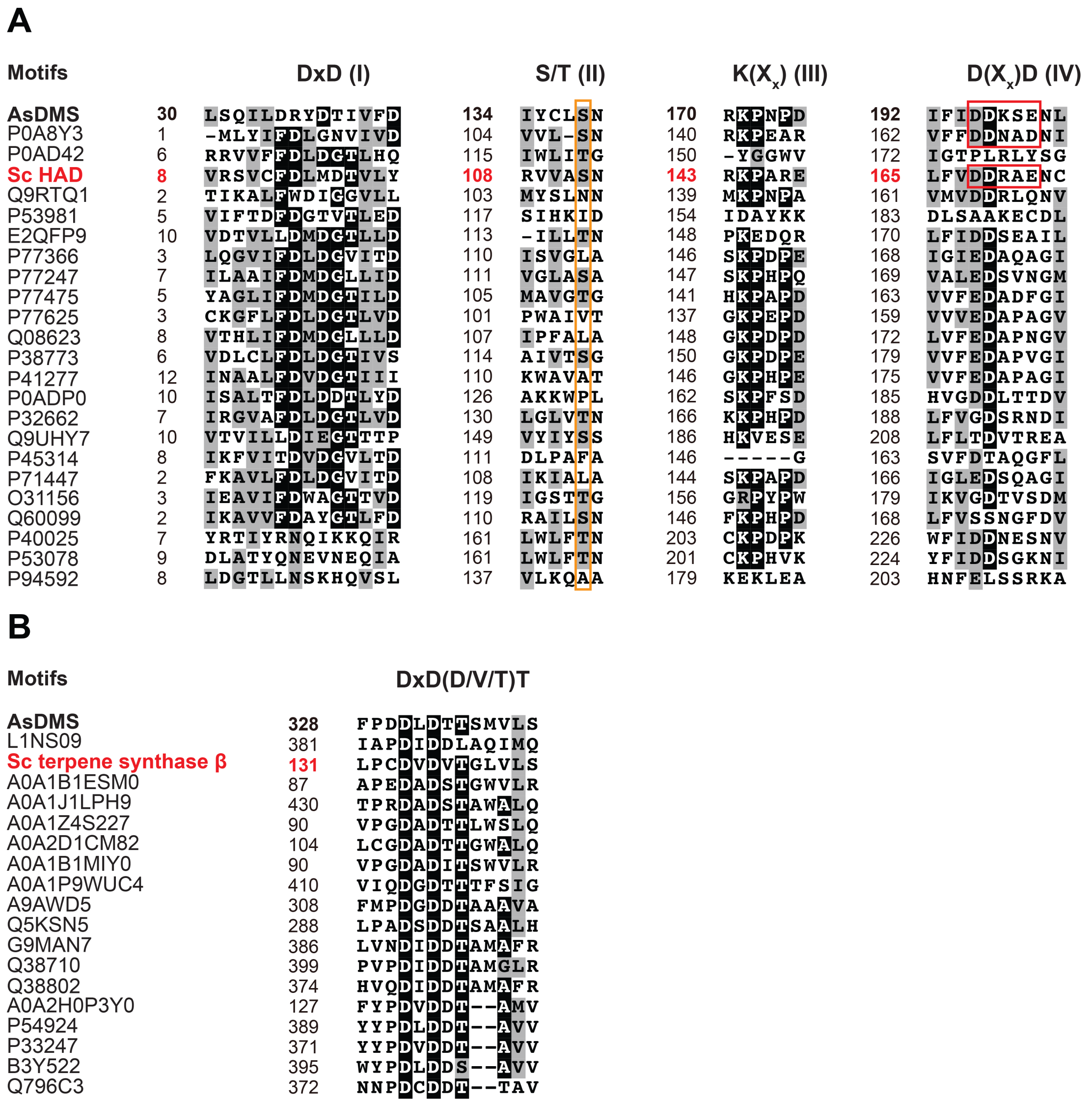


**Figure S10.** Sequence comparisons of HAD and terpene synthase proteins in *S. cellulosum* with selected HAD-like hydrolases and β domains of terpene synthases (accession numbers provided), respectively. (A) Sequence alignment of Sc HAD (red bold text) with the N-domain of AsDMS (black bold text) and selected HAD-like hydrolases highlights four conserved HAD signature motifs: DxD (motif I), S/T (II), K($X_x$) (III), and D($X_x$)D (IV). Red boxes highlight the typical DDxxE or DDxxD motifs of terpene synthases. Orange box indicates the position of the S/T motif. (B) Comparisons of Sc terpene synthase β (red bold font) with the C-domain of AsDMS (black bold text) and selected β domains of terpene synthases indicate the presence of the DxDVT motif in Sc terpene synthase β that corresponds closely to the DxDTT motif in AsDMS or DxDDT/DxDTT motifs of other enzymes. Protein sequences were aligned and colored using GenomeNet ClustalW v1.83 and the BoxShade 3.21 server, respectively. As, *Aquimarina spongiae*; Sc, *Sorangium cellulosum*.

**A**

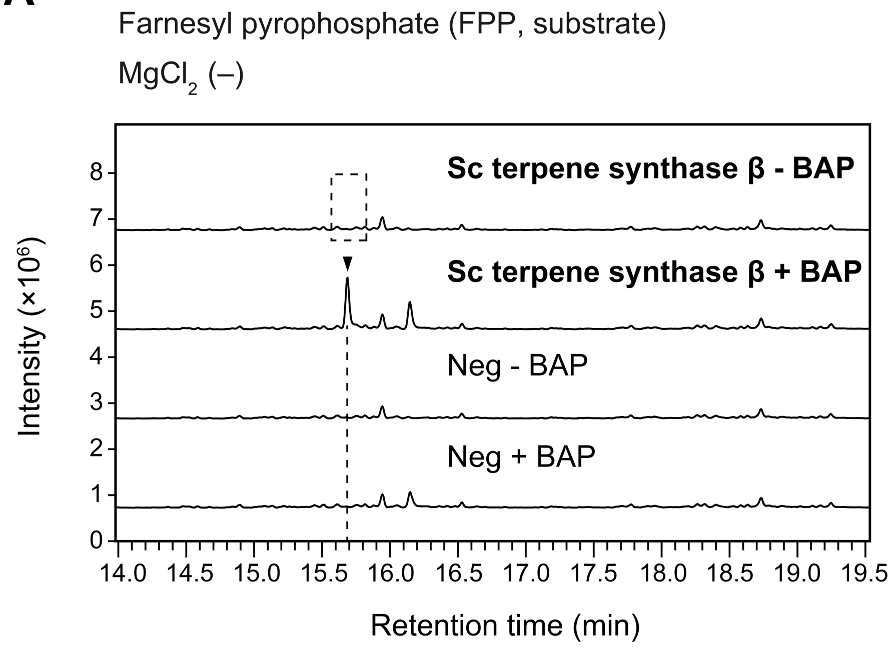


**B**

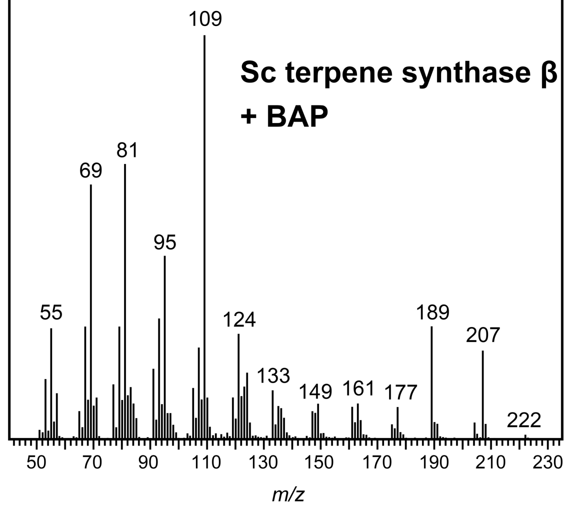


**Figure S11.** *In vitro* enzymatic assays of Sc terpene synthase β protein in the absence of $Mg^{2+}$. (A) GC-MS chromatograms of *in vitro* reaction products. Enzymatic assays of purified Sc terpene synthase β protein with the FPP substrate were performed individually in the absence of $MgCl_2$ with or without alkaline phosphatase (BAP) treatment. Reactions in the absence of protein were used as negative controls (Neg). Data represent three independent reactions performed with the same preparation of purified protein. Dashed boxes indicate that no desired reaction products were detected. (B) Mass spectrum of the reaction product detected in (A). Sc, *Sorangium cellulosum*. *m/z*, mass-to-charge ratio.

**Method S1**

**Expression, Purification, and *In Vitro* Enzymatic Activity Assays of Putative ABM Proteins.** All recombinant N-terminal $His_8$-tagged ABM proteins were expressed in *E. coli* BL21 Star (DE3) strains harboring pET28b(+)-$His_8$*ABM*. The purification of these proteins was performed as described for recombinant FPPS proteins in this study followed by *in vitro* enzymatic activity assays. Several reaction conditions were tested: a total volume of 200 $\mu$L of reaction mixture containing 50 mM Tris-HCl (pH 8.0), 2 mM $MgCl_2$, 100 $\mu$M FPP or farnesol or drimenol, 100 $\mu$M NADPH, 100 $\mu$M FAD, and 1 $\mu$M purified ABM proteins at 30 °C within 1, 4, or 18 h. The extraction was performed twice using 200 $\mu$L of hexane/ethyl acetate (1:1, v/v) and further centrifuged at 4,400 × *g* for 5 min. The combined organic layer was subjected to GC-MS analysis, as described for the characterization of FPPSs.

**Method S2. Materials for Cloning and Enzyme Activity Assays.**

**Bacterial Strains and Plasmids.** The *E. coli* HST08 strain was purchased from Takara Bio. Plasmid pET28b(+) and *E. coli* BL21 Star (DE3), which were used for heterologous expression of FPPS, Sc HAD, or Sc terpene synthase β proteins, were purchased from Merck Millipore (Burlington, MA, USA) and Thermo Fisher Scientific (Waltham, MA, USA), respectively.

**Chemicals and Enzymes.** Substrates, authentic samples, and cofactors used for enzymatic assays and GC-MS analysis, including FPP, farnesol, NADPH, and FAD were obtained from Sigma-Aldrich (St. Louis, MO, USA). Authentic (–)-drimenol was supplied from Cayman Chemical (Ann Arbor, MI, USA). We used (+)-nootkatone (Sigma-Aldrich) as the internal standard. Magnesium chloride and dithiothreitol were obtained from Nacalai Tesque (Kyoto, Japan). All organic solvents (Wako Pure Chemical Industries, Osaka, Japan) were of analytical grade. Purified water was prepared using the Milli-Q purification system (Millipore, Billerica, MA, USA). Restriction enzymes and BAP from *E. coli* C75 were purchased from Takara Bio. Pre-cast SDS-PAGE gels (Wako Pure Chemical Industries) were used for protein expression and purity analyses. Coomassie Brilliant Blue (EzStain Aqua) obtained from ATTO (Tokyo, Japan) was used for gel staining.
